# Parvalbumin interneuron dendrites enhance gamma oscillations

**DOI:** 10.1101/2021.06.22.449483

**Authors:** Birgit Kriener, Hua Hu, Koen Vervaeke

## Abstract

Dendrites are important determinants of the input-output relationship of single neurons, but their role in network computations is not well understood. Here, we used a combination of dendritic patch-clamp recordings and in silico modeling to determine how dendrites of parvalbumin (PV)-expressing basket cells contribute to network oscillations in the gamma frequency band. Simultaneous soma-dendrite recordings from PV basket cells in the dentate gyrus revealed that the slope, or gain, of the dendritic input-output relationship is exceptionally low, thereby reducing the cell’s sensitivity to changes in its input. By simulating gamma oscillations in detailed network models, we demonstrate that the low gain is key to increase spike synchrony in PV neuron assemblies when cells are driven by spatially and temporally heterogeneous synaptic input. These results highlight the role of dendritic computations in synchronized network oscillations.

## Introduction

Network oscillations in the gamma frequency band (40-110 Hz) are a prominent circuit feature of many brain areas and likely support a variety of cognitive processes such as perception (Gray et al., 1989), attentional selection (Fries et al., 2001), and memory (Lisman and Idiart, 1995; Lundqvist et al., 2016). The important roles for gamma oscillations have triggered numerous studies investigating how they are generated. These show that parvalbumin-expressing inhibitory neurons, that form axonal “baskets” around the soma of target neurons (hence called PV basket cells (Hu and Jonas, 2014)), play a central role in the cortex (Fuchs et al., 2007; Cardin et al., 2009; Sohal et al., 2009). Although previous studies have identified a number of synaptic mechanisms in basket cells that seem optimal to generate gamma oscillations (Bartos et al., 2002, 2001; Vida et al., 2006; Strüber et al., 2015; Bartos et al., 2007; Erisir et al., 1999; Pike et al., 2000; Hormuzdi et al., 2001; Whittington et al., 1995; Fisahn et al., 1998; Cornford et al., 2019; Kopell and Ermentrout, 2004), the dendritic properties that allow PV cells to transform synaptic inputs into synchronous output firing remain elusive.

Spike synchrony in an ensemble of inhibitory interneurons is a key mechanism for generating network oscillations (Bartos et al., 2007; Buzsáki and Wang, 2012). In the simplest of models, when a sufficiently large number of inhibitory neurons fires within in a small time window, it generates an pronounced inhibitory conductance in the network. If the excitatory input drive is homogeneous across cells, neurons will escape inhibition together and fire subsequently at the same time, leading to synchronous activity (Bartos et al., 2007; Buzsáki and Wang, 2012). However, when the excitatory drive varies from neuron to neuron, cells will fire at different rates, escape the common rhythm, and synchrony is lost (Wang and Buzsáki, 1996; Wang, 2010). In biological networks, a plethora of heterogeneities enhance spike rate variability (Softky and Koch, 1993). Each cell, even of the same type, has a different excitability and morphology, and spatially distributed cells receive different amounts of excitation (spatial heterogeneity) which also fluctuates rapidly over time (Destexhe et al., 2003; Calvin and Stevens, 1967) (temporal heterogeneity). These different forms of heterogeneity have been a long-standing challenge of network models studying synchrony (Bartos et al., 2002, 2001; Vida et al., 2006; Bartos et al., 2007; Wang and Buzsáki, 1996; Moca et al., 2012; Tikidji-Hamburyan et al., 2015; Tiesinga and José, 2000; Tort et al., 2007; Neltner et al., 2000; White et al., 1998).

The unique biophysical properties of PV basket cells further exacerbate this problem. How neurons integrate sustained excitatory synaptic input and transform it into output firing rate is captured by their input-output (I-O) relationship (Silver, 2010). This is typically measured by injecting current in the soma and measuring spike frequency. In the case of PV basket cells, the gain of the I-O relationship is about ten times steeper compared to cortical principal neurons (Goldberg et al., 2008). Therefore, in an ensemble of PV neurons that all receive different amounts of input, the cells will spike at dramatically different rates, making it challenging to synchronize (illustrated in Fig. 1a).

**Figure 1.**
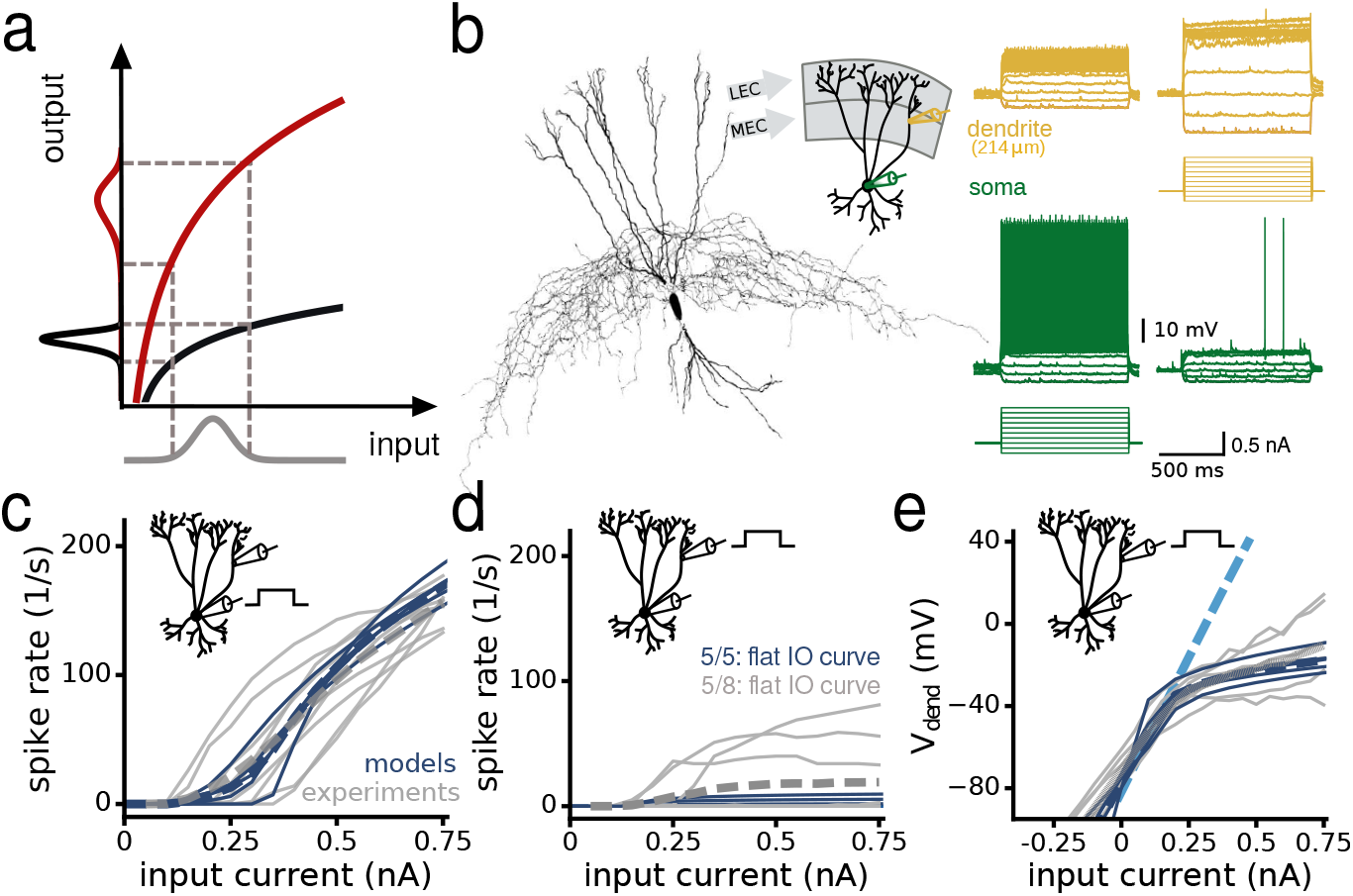
Dendritic input results in a lower I-O gain compared to somatic input. **(a)** A steeper hypothetical input-output (I-O) relationship maps the same input distribution to a broader output firing rate distribution. **(b)** Example of a dual soma-dendritic patch-clamp recording of a parvalbumin-expressing (PV) basket cell in the dentate gyrus. Cartoon shows the recording configuration and the input from the lateral and the medial entorhinal cortex (LEC, MEC). Membrane potential traces show soma and dendrite responses to current injections in either the soma (left) or the dendrite (right). Inset, example of a reconstructed morphology of a PV basket cell used for computer simulations. **(c)** I-O relationships when injecting current steps in the soma. Grey lines, individual experiments (n=8); Blue lines, individual models (n=5); Dashed thick lines, averages. **(d)** I-O relationships when injecting current steps in the dendrite. Average dendritic recording distance is 240 ± 7 μm (n = 8 experiments) and 230 μm from the soma (n=5 models). In 5/8 experiments and 5/5 models, cells fired only a few or no spikes, resulting in near flat I-O relationships. **(e)** Dendritic membrane potential in response to current steps injected in the dendrite (n = 6 experiments, n = 5 models). Light blue dashed line is the average dendritic membrane potential when no voltage-dependent K^+^-channels are present on the dendrites.

Previous work on gamma oscillations only performed network simulations using so-called “point-neuron” models that disregard the dendritic morphology. However, we hypothesized that PV basket cell dendrites are key for enhancing the robustness of gamma oscillations in heterogeneous networks. We made simultaneous whole-cell recordings from the dendrites and soma of PV cells in the dentate gyrus of the rat hippocampus, a circuit that generates prominent gamma oscillations (Towers et al., 2002; Csicsvari et al., 2003; Hu et al., 2010). We found that, compared to the steep I-O relationship measured from the soma, the gain of the dendritic I-O relationship is scaled down. Therefore, PV cells are far less sensitive to different amounts of input than previously thought (Wang and Buzsáki, 1996). Furthermore, PV cell dendrites reduce the amplitude of fast-fluctuating synaptic input, enhancing regular spiking. We used anatomically detailed networks models to show that PV cell dendrites indeed dramatically increase the robustness of gamma oscillations in a wide variety of network architectures. Closer examination of the underlying biophysical mechanisms revealed that the high-threshold and fast activating K^+^-currents in the dendrites (Hu et al., 2010) act to dampen heterogeneities and thereby enhance spike synchrony.

## Results

### Dendritic input results in a lower I-O gain compared to somatic input

To compare the I-O relationship of soma-driven and dendrite-driven output firing in PV basket cells, we made dual whole-cell recordings from the soma and dendrites using confocal microscope-guided patching (Hu et al., 2010). We targeted the dendrites in the middle and outer third of the molecular layer where these cells receive input from the medial and lateral entorhinal cortex, respectively (Amaral et al., 2007) (Fig. 1b, range: 214 to 262 μm from the soma; 240 ± 7 μm, n = 8 cells). Current injections of increasing amplitude in the soma increased the spike frequency rapidly, leading to a steep I-O relationship (Fig. 1c, gain = 390 ± 26 Hz nA^-1^). In stark contrast, current injection in the dendrites triggered no, or low-frequency, spiking, resulting in a nearly flat average I-O relationship (Fig. 1d). Inspection of the dendritic membrane potential revealed that responses to local current injection were linear up to ~ −30 mV but then became sub-linear, making it increasingly hard to drive the cell to fire (Fig. 1e, and example in b). These data show that, compared to the soma, the gain of the dendritic I-O relationship is scaled down.

Previous work has characterized the biophysical properties of PV neurons, such as the passive electrical membrane properties and ion channel distributions (Hu et al., 2014, 2010; Nörenberg et al., 2010; Hu and Jonas, 2014). To test whether these known properties can account for the low I-O gain, we used a computational approach. We simulated the I-O relationship using five anatomically detailed PV neuron models that included dendrites and the axon (example in Fig.1b, Suppl. Fig. S1a, see Methods PV Basket Cell Model). The only voltage-dependent conductance we inserted in the dendrites was a high-threshold and fast-activating K^+^-channel that is a hallmark of PV basket cells in the dentate gyrus (Hu et al., 2010). These models recapitulated the fast-spiking phenotype, the strong attenuation of backpropagating action potentials (Hu et al., 2010) and the steep soma I-O relationship (Fig. 1c, Suppl. Fig. S1b,c, gain = 370 ± 32 Hz nA^-1^, n = 5 models). Consistent with the majority of experiments, injecting current into the distal dendrites produced a nearly flat I-O relationship (Fig. 1d, Suppl. Fig. S1b,c, distance = 230 μm from the soma). Furthermore, the models recapitulated the sub-linear increase of the dendritic membrane potential, which was dependent on the dendritic K^+^-channels (Fig. 1e).

In summary, the PV basket cell models capture the essential I-O properties that we measured experimentally, and collectively, these data show that the dendrite-driven I-O relationship has a lower gain compared to soma-driven output firing.

### Distributed dendritic input results in a lower I-O gain and more regular spiking

In vivo, PV basket cells likely receive both clustered and spatially distributed input on the dendritic tree. Dual soma-dendritic patch-clamp recordings, however, can only mimic clustered dendritic input. Therefore, to examine how distributed input affects the IO relationship, we used the PV neuron models. To perform these simulations, we needed to verify that the models could reproduce some of the most elementary properties of dendritic integration such as the kinetics and the attenuation of excitatory synaptic potentials (EPSPs) along the dendrites. Therefore, we further analyzed the soma-dendritic experiments. The amplitude of dendritic EPSPs increased with distance from the soma, while their amplitude at the soma decreased (Fig. 2a, n = 5 cells). The space-constant of dendrite-to-soma attenuation was 84 μm, and this was similar to the value predicted by our models (74 ± 19 μm, n = 5 models, Fig. 2b).

**Figure 2.**
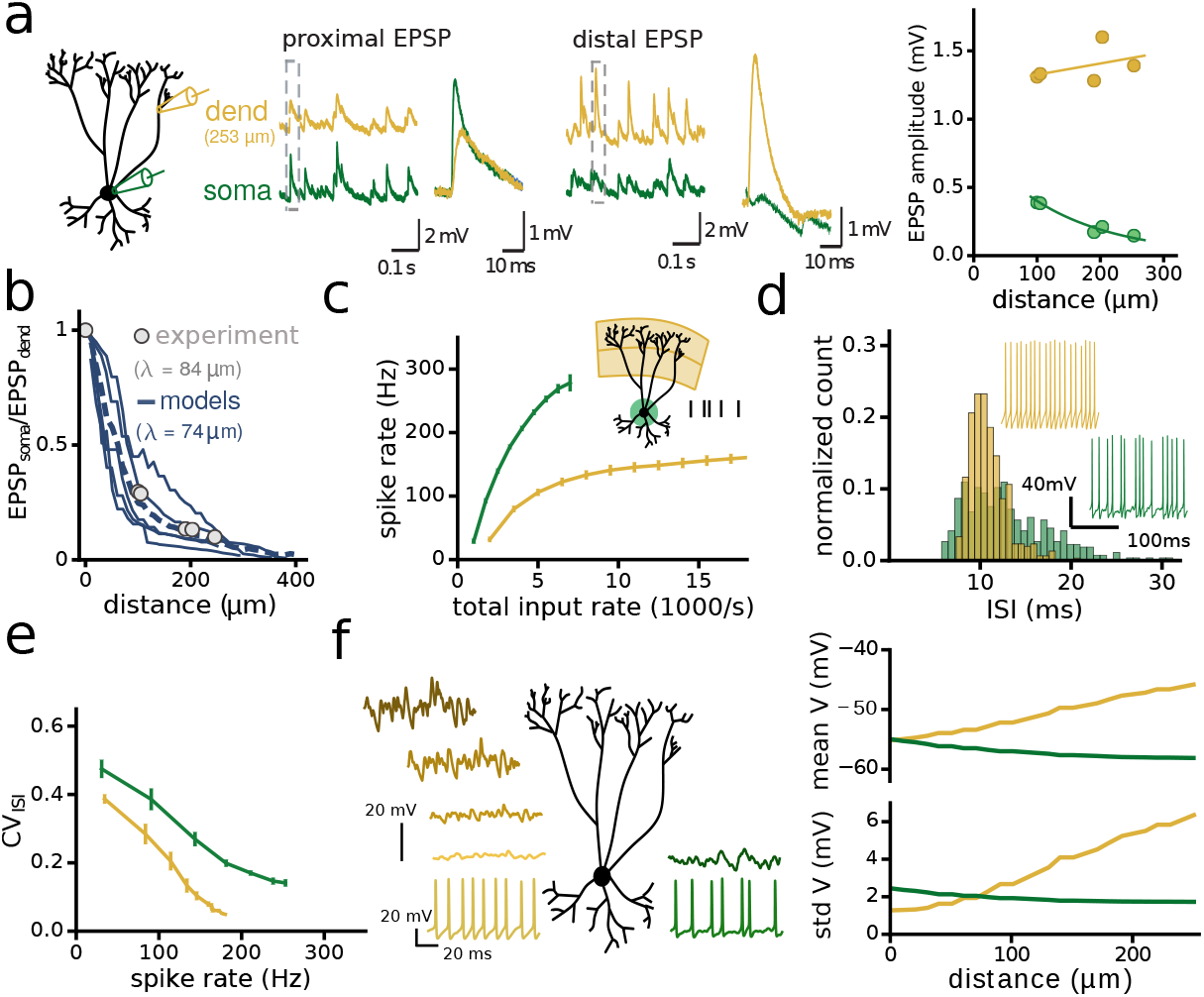
Distributed dendritic input results in a lower I-O gain and more regular spiking. **(a)** Cartoon shows the recording configuration. Membrane potential traces show spontaneous EPSPs. We used the fast-rising slope (<0.5 ms) to determine which EPSPs originate close to the recording pipette. Right panel shows the mean EPSP amplitude of all cells, plotted as a function of input distance from the soma, for both dendritic (orange) and somatic recordings (green, n=5). **(b)** EPSP amplitude attenuation during propagation from the dendrites to the soma. Grey data points, experiments (n=5); Blue lines, model predictions (n=5); Blue dashed line, average model prediction. Data and average model prediction were fitted with a single exponential decay function (~ *e^-x/λ^* with length constant *λ*). **(c)** I-O relationships of PV cell models using perisomatic (green) or distributed dendritic input (orange, n=5 models). Cartoon shows the synaptic input locations; perisomatic: 50 synapses ≤ 50 μm from the soma. Dendritic: 100 synapses ≥ 120 μm from the soma. Black ticks illustrate a Poisson train of synaptic inputs. Synapses were randomly distributed. Total input rate is the number of synapses × the rate per synapse. The number of synapses was fixed, and we varied only the rate per synapse. **(d)** Insets, example spike trains from the data in (c). Normalized count histograms show the interspike intervals (ISI) of the example spike trains. **(e)** ISI irregularity quantified by the Coefficient of Variation (CV) as a function of output spike rate (based on the data in c). **(f)** Membrane potential traces along the dendrites and the soma, when the input is dendritic- (orange) or perisomatic (green). For dendritic input, membrane potential fluctuations attenuate towards the soma, leading to more regular spiking. Right panel shows the mean and standard deviation of the membrane potential as a function of distance to soma. The amount of input was adjusted to achieve the same mean depolarization of the soma. Na+ channels were blocked to prevent action potentials.

Having validated the PV neuron models, we next simulated the I-O relationship for distributed synaptic input. We randomly positioned excitatory synapses on the outer two-thirds of the dendrites (≥ 120 μm from the soma) or within 50 μm from the soma (Fig. 2c). Simulating sustained input revealed that the gain of the I-O relationship for dendritic drive was again lower than for somatic drive (Fig. 2c).

In addition, we observed that the dendrite-driven output firing was also more regular, as illustrated by the narrower distribution of spike intervals (Fig. 2d), a property that may help spike synchronization between PV neurons. Quantifying the spike interval variability by the coefficient of variation (CV, the ratio of the standard deviation to the mean) showed that dendrite-driven firing was more regular across the entire spike frequency range (Fig. 2e). We hypothesized that this was due to the strong EPSP attenuation along the dendrites (Fig. 2b); Because distal dendritic EPSPs are small when arriving at the soma compared to proximal EPSPs (Fig. 2a), the membrane potential fluctuations are also smaller, leading to more regular spiking. To test this, we examined the standard deviation of the membrane potential along the dendrites during dendritic stimulation (Fig. 2f). These data show that, indeed, during sustained dendritic input, the membrane potential fluctuations are small near the soma.

In summary, these data suggest that PV basket cell dendrites may enhance gamma synchrony in two ways. First, the low gain of the dendritic I-O relationship will reduce the sensitivity of PV basket cell output to different amounts of input, and second, more regular spiking may facilitate spike synchronization between PV cells.

### PV neuron dendrites make gamma synchrony more robust to heterogeneities

To test whether PV cell dendrites enhance gamma synchrony in heterogeneous networks, we performed network simulations using the reconstructed PV cells coupled with inhibitory synapses based on empirical data (Bartos et al., 2002) (Fig. 3a, see Methods Network models). We included two forms of heterogeneity; (1) Each PV cell received a different amount of synaptic input to create spatial heterogeneity (so-called “input heterogeneity”), and (2), the input consisted of noisy Poisson trains of synaptic conductances to create temporal heterogeneity. Each cell in the network received a mean input rate taken from a normal distribution with a mean μ and standard deviation δ. We could then increase the spatial heterogeneity in the network by increasing the width of this distribution (that is, the ratio of *μ/δ*×100). In homogenous networks (0% input heterogeneity), spike synchrony in the gamma frequency range emerged rapidly in both soma-driven and dendrite-driven networks (Fig. 3b). However, when we increased the input heterogeneity, dendrite-driven neurons fired more synchronously (Fig. 3b, compare raster plots).

**Figure 3.**
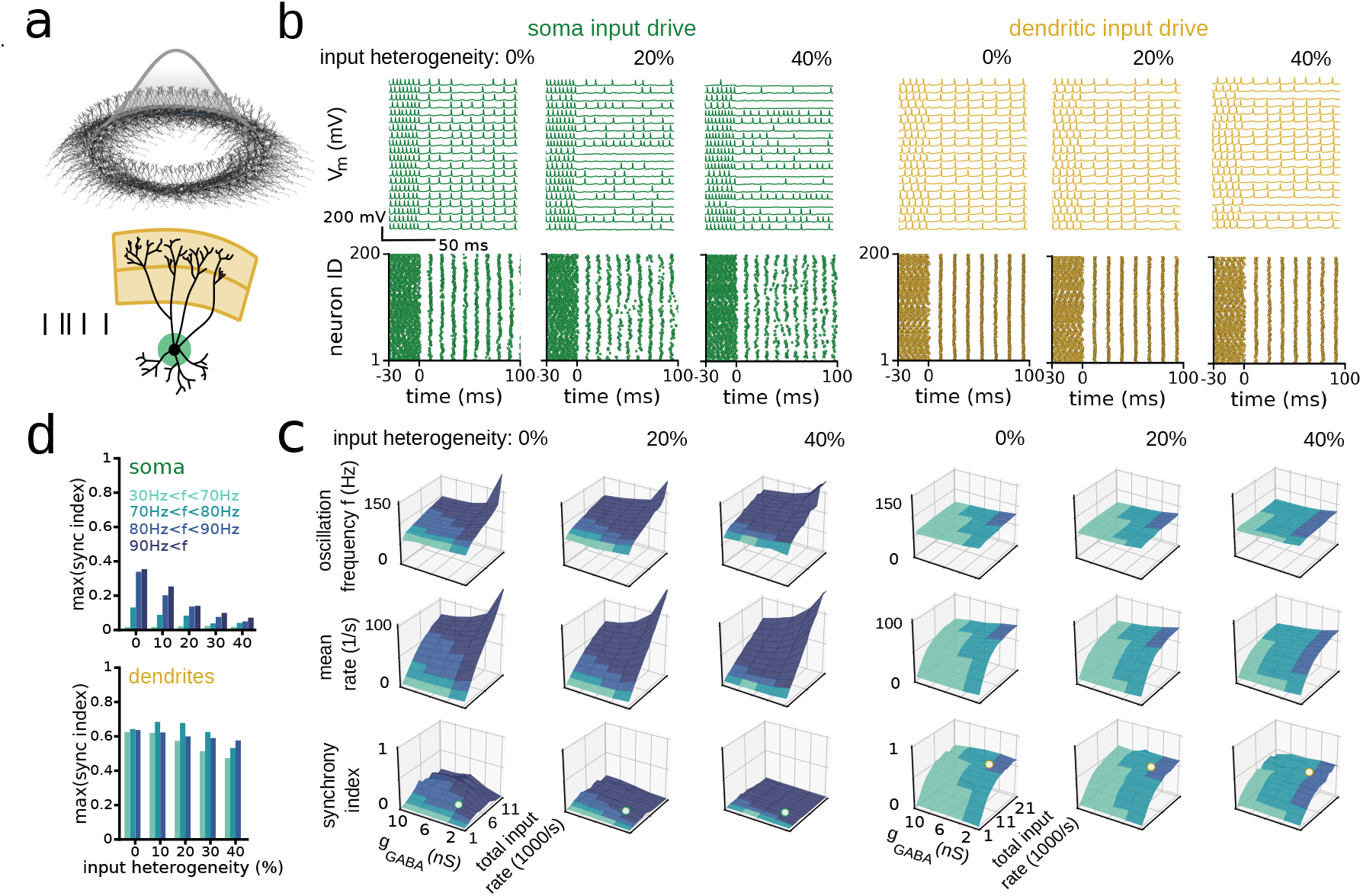
PV neuron dendrites make gamma synchrony more robust to input heterogeneity. **(a)** Ring network of 200 PV neuron models with a reconstructed morphology, symbolically arranged along a ring to illustrate a network with local inhibitory connections. Cells are randomly coupled by inhibitory synapses following a Gaussian connection probability (grey curve). Cartoon shows the synaptic input locations; 50 synapses ≤ 50 μm from the soma (green); 100 synapses ≥ 120 μm from soma (orange). **(b)** PV cell activity in networks driven by perisomatic (green) or dendritic (orange) excitation. Input heterogeneity increases from left to right. Top row, example membrane potential traces showing spikes from 20 random cells. Bottom row, raster plots of all 200 PV cells in the network. Network starts uncoupled and inhibitory synapses activate at t = 0 ms. **(c)** Network oscillation frequency (top row), mean spike rate (middle row) and Synchrony Index (bottom row) as a function of total input rate and the strength of inhibitory conductance (gGABA). The surface colours show the oscillation frequency range (see colour legend in d). White dots on the Synchrony Index corresponds to the examples in (b). **(d)** The maximum Synchrony Index per frequency band, as a function of input heterogeneity for perisomatic and dendritic input (based on data in (c)).

The amplitude of the excitatory drive and the strength of inhibitory connections are critical determinants of the spike rates, oscillation frequency and synchrony, therefore, we examined how they affect the results (Fig. 3c). To quantify network synchrony, we defined the Synchrony Index, which is 1 for perfect spike synchrony and approaches 0 when the network is fully desynchronized (see Methods Analysis). For soma-driven networks, increasing the input heterogeneity to 40% reduced the Synchrony Index for a broad range of excitation and connection strengths (Fig. 3c, bottom row). In stark contrast, the dendrite-driven networks maintained a high Synchrony Index (Fig.3c). To summarize the relationship between Synchrony Index and heterogeneity for different gamma frequency bands, we divided the gamma spectrum into smaller frequency bands and calculated the maximum Synchrony Index for each band (Fig. 3d). These data show that regardless of the gamma frequency range, dendrite-driven networks are far more tolerant to heterogeneities. Even increasing the input heterogeneity to 100% caused only a small reduction in network synchrony (Suppl. Fig. S2a). Furthermore, dendrite-driven networks synchronized robustly even when the strength of inhibitory connections was an order of magnitude less compared to previous influential models (Vida et al., 2006) (see side-by-side comparison in Suppl. Fig. S2a-c). Finally, we found similar results regardless of which PV basket cell model we used for building networks, illustrating that the results do not depend on a specific cell morphology or specific biophysical properties (Suppl. Fig. S2d).

Notably, we observed that network synchrony in dendrite-driven networks was also higher in homogeneous networks in which all cells receive the same input (Fig. 3d, 0% input heterogeneity case). Therefore, we hypothesized that the dendrites also reduced temporal heterogeneities by reducing the amplitude of the membrane potential fluctuations at the soma (see Fig. 2f). This could facilitate spike synchrony by enhancing spike regularity (see Fig. 2d,e). To test this, we substituted the Poisson synaptic inputs with tonic input currents to produce noiseless excitation (Suppl. Fig. S3). Indeed, synchrony in soma-driven and dendrite-driven networks was now similar (Suppl. Fig. S3d, 0% input heterogeneity case). Altogether these data show that both spatial and temporal heterogeneities are important determinants of network synchrony, and that dendrite-driven synchrony is more robust to both for a wide range of parameters.

So far, we used the most elementary network models that produce gamma synchrony. To investigate whether our results depend on specific neuronal or network properties, we performed additional simulations. *First*, we explored the impact of adding electrical synapses (Hormuzdi et al., 2001) (Suppl. Fig. S4a-c). Electrical synapses between PV neurons increase the Synchrony Index for both dendrite-driven and soma-driven networks but do not change the conclusion that dendrite-driven synchrony is more robust (Suppl. Fig. S4c). *Second*, adding N-Metyl-D-Aspartate (NMDA) receptors (Koh et al., 1995b; Sambandan et al., 2010) enhances the I-O gain for both somatic and dendritic input (Suppl. Fig. S4d), but did also not change our conclusion (Suppl. Fig. S4f). *Third*, because inhibition can be hyperpolarizing or shunting (Vida et al., 2006), we performed network simulations while varying the reversal potential of inhibition. However, this did also not affect the outcome (Suppl. Fig. S4g-i). *Fourth*, when we changed the network from a 1D “ring structure” to a 2D network with local connectivity based on recent empirical data (Espinoza et al., 2018), dendrite-driven gamma synchrony was also more robust (Suppl. Fig. S5a-c). Finally, we constructed composite networks by using a mixture of all five reconstructed PV neuron models. Because the models have a substantially different excitability (input resistance varies between 57-119MΩ) and morphology (Suppl. Fig. S1a), this cell-to-cell variability adds further spatial heterogeneity and increases the realism of the network. Nevertheless, dendrite-driven gamma synchrony remained more robust (Suppl. Fig. S5d-f).

In summary, using anatomically and biophysically detailed network models, we found that input arriving on the dendrites strengthens the robustness of gamma synchrony in heterogeneous networks, and we show that these results hold for a wide range of neuronal and network parameters.

### How do PV basket cell dendrites enhance gamma oscillations?

This question is difficult to address in anatomically and biophysically detailed PV neuron models because manipulating the dendritic properties also affects the fast-spiking phenotype (Hu et al., 2010). Therefore, we constructed a simpler model to determine the underlying biophysical mechanisms. We used a single soma compartment and five apical dendrites with similar length and diameter as real PV basket cells (Fig. 4a). For the spiking mechanism, we used a well-known model that recapitulates the spiking properties of these cells (Wang-Buzsáki model (Wang and Buzsáki, 1996)), and we added the high-threshold activated K^+^-conductance also to the dendrites (Hu et al., 2010), see Methods Ball-and-sticks model. The model resembled the measured I-O properties (Suppl. Fig. S6a), and reproduced the Synchrony Index of the anatomically-detailed models (Fig. 4a,b, Suppl. Fig. S6b-e).

**Figure 4.**
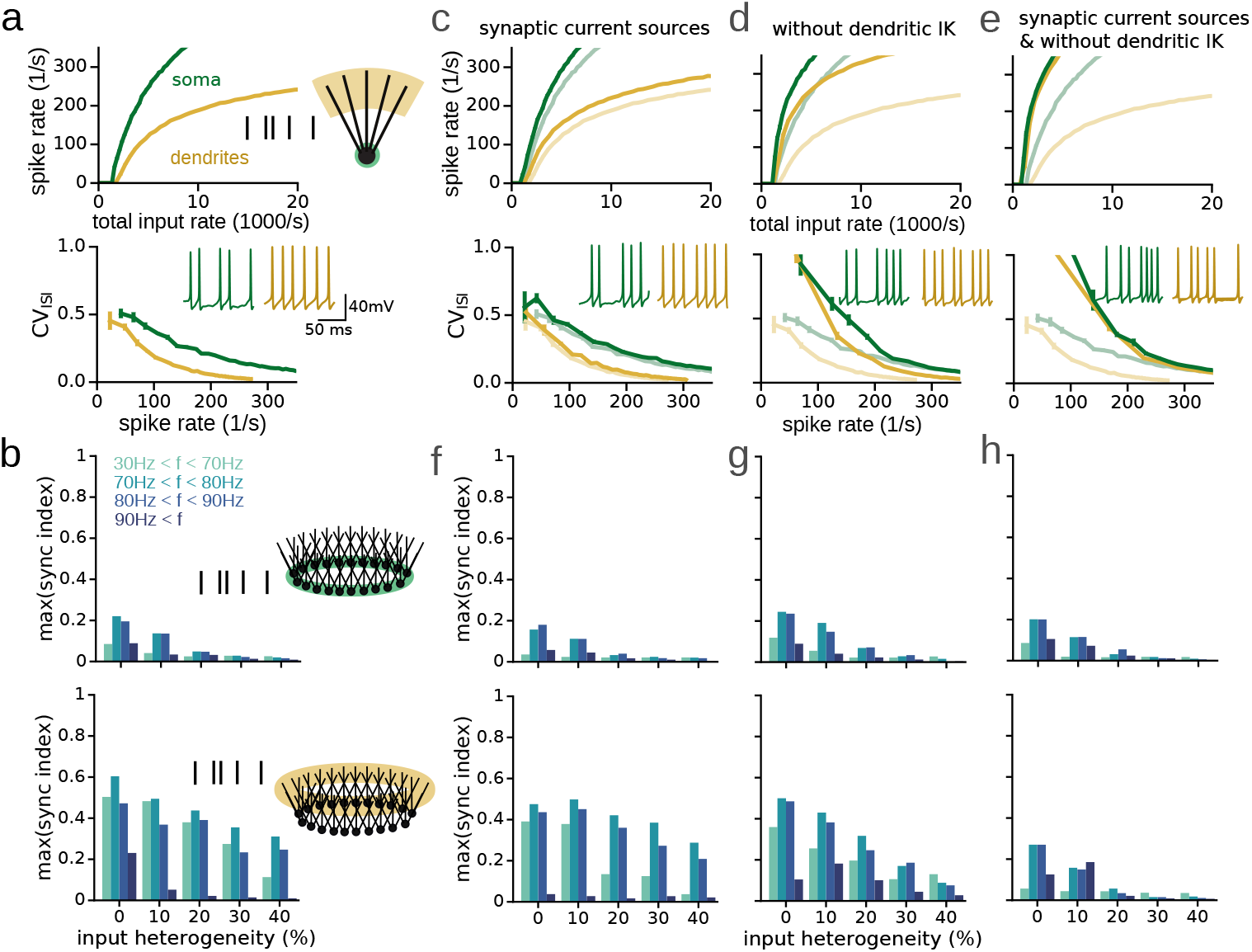
Biophysical mechanisms of PV neuron dendrites that underlie robust gamma oscillations. **(a)** Top, I-O relationships of a simplified PV neuron model (cartoon), driven by excitatory input on the soma (green, 50 synapses) or dendrites (orange, 100 synapses, ≥ 150 μm from the soma). Bottom, coefficient of variation (CV) of the interspike intervals (ISI) as a function of output spike rate. Insets, example spike trains. **(b)** Data from ring networks using 200 simplified PV cell models, driven by either soma or dendritic input. Histograms show maximum Synchrony Index per frequency band, as a function of input heterogeneity. Colour legend indicates the oscillation frequency bands. **(c)** As (a), but using synaptic current sources instead of synaptic conductances (these synapses have the same kinetics, but with a fixed driving force). Faint lines are the data from (a) for comparison. **(d)** As (a), but without dendritic voltage-dependent K^+^-channels. **(e)** As (a), but using synaptic current sources and without dendritic K+ channels. **(f,h)** The maximum Synchrony Index per frequency band, as a function of input heterogeneity, based on ring networks built from PV models described in (c-e). The three different conditions can be compared with the control histograms in (b).

We explored two hypotheses to explain the robustness of dendrite-driven synchrony. First, thin PV cell dendrites have a high impedance and strongly depolarize when excited by synaptic inputs. Therefore, a reduced driving force in the dendrites will limit the synaptic current generated (Bush and Sejnowski, 1994) and will decrease the I-O gain. A reduced driving force will also reduce the amplitude of the membrane potential fluctuations and enhance spike regularity. To test this hypothesis, we converted excitatory synaptic conductances to currents (which are independent of driving force). Synaptic currents indeed increased the gain of the I-O relationship (Fig. 4c) but did not significantly affect spike regularity (quantified by the CV), and the Synchrony Index was only slightly reduced (compare Fig. 4f with Fig. 4b). Therefore, this mechanism appears to play only a minor role in enhancing network synchrony, and other mechanisms must exist.

A second hypothesis is that the K^+^-conductances in the dendrites reduce the gain of the dendritic I-O relationship by actively opposing dendritic depolarization (Hu et al., 2010). Deleting the K^+^-conductance from the dendrites indeed increased the gain (Fig. 4d) and spike intervals became more irregular (Fig.4d). Furthermore, dendrite-driven synchrony became slightly more sensitive to input heterogeneity (Fig. 4g). However, the difference between soma-driven and dendrite-driven synchrony was still striking, indicating that the K+ conductance alone does not explain why dendrite-driven synchrony is more robust (Fig. 4g).

Finally, we considered the following possibility. We hypothesized that the depolarization caused by deleting the voltage-dependent K^+^-conductances –which should steeply increase the gain– may be limited because it is compensated by a strong reduction in driving force for synaptic excitation. To test this, we deleted the K^+^-conductance from the dendrites and we used synaptic currents. In this condition, the gain of both the soma-driven and dendrite-driven I-O relationship sharply increased (Fig. 4e). Furthermore, spike intervals became very irregular (Fig.4e). And finally, dendrite-driven networks were now equally sensitive to input heterogeneity as the soma-driven networks (Fig. 4h).

Altogether, these data show that dendritic K^+^-currents, and to a lesser degree, the reduced driving force for excitation, decrease the I-O gain (enhancing robustness to input heterogeneity) and reduce membrane potential fluctuations (enhancing robustness to temporal heterogeneity).

### PV dendrites enhance the robustness of theta-nested gamma rhythms

Gamma oscillations often occur superimposed on the slower theta (5-12 Hz) oscillations (Bragin et al., 1995; Pernía-Andrade and Jonas, 2014). Such cross-frequency coupling may serve to couple remote cortical circuits (Colgin, 2015). Therefore, we tested whether dendrites also enhance gamma synchrony when modulated by the theta rhythm. We used the anatomically detailed network models and simulated the theta rhythm by a sinusoidal current (Fig. 5a, see Methods External drive). We then compared the robustness of gamma synchrony when driving the network with somatic or dendritic input.

**Figure 5.**
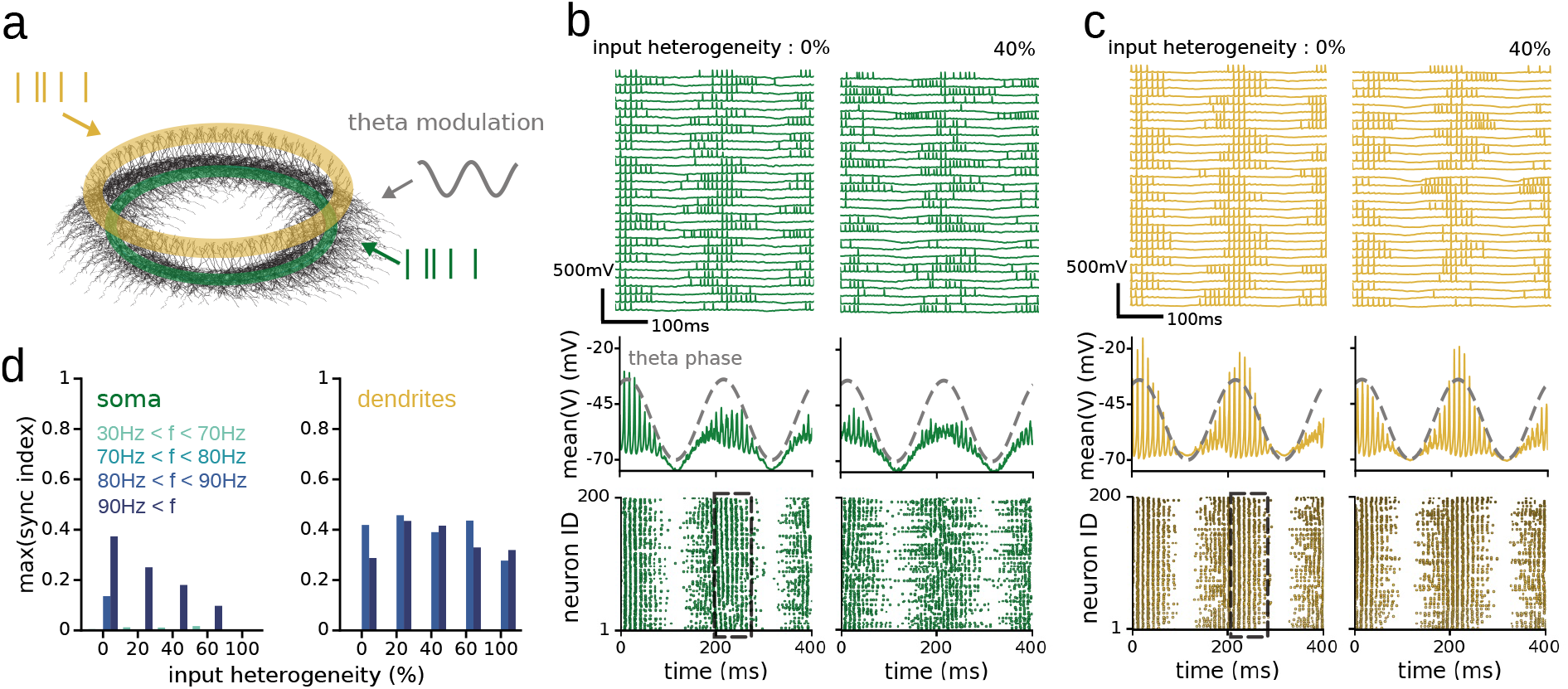
PV dendrites enhance the robustness of theta-nested gamma rhythms. **(a)** Ring network of 200 PV neuron models with a reconstructed morphology. PV cells receive a theta frequency-modulated input current to the soma. In addition, networks receive rate-coded excitatory input either close to the soma (green, 50 synapses ≤ 50 μm from the soma) or to the dendrites (orange, 100 synapses ≥ 120 μm from soma). **(b)** PV cell activity in a network driven by perisomatic excitation. Input heterogeneity increases from left to right. Top row, example membrane potential traces showing spikes from 30 random cells. Middle row, mean membrane potential. Dashed line is the theta phase. Bottom row, raster plots of all 200 PV cells in the network. Black box, phase-interval during which the Synchrony Index was calculated. **(c)** As (b) but networks are driven with dendritic excitation. **(d)** The maximum Synchrony Index per frequency band, as a function of input heterogeneity.

Similar to observations in the dentate gyrus of exploring rats (Bragin et al., 1995), the simulated gamma frequency was in the range of 80-100 Hz. For homogeneous networks (0% heterogeneity), both soma and dendrite-driven synchrony emerged within a single theta cycle (Fig. 5b,c). Furthermore, dendrite-driven synchrony was substantially higher compared to soma-driven synchrony (Fig. 5b,c). With increasing input heterogeneity, soma-driven network synchrony fell rapidly, while dendritic-driven synchrony remained intact for all frequency bands (Fig. 5d). These data show that dendrite-driven gamma synchrony superimposed on theta oscillations is also more robust to network heterogeneities.

### PV dendrites enhance synchrony in networks of synaptically coupled excitatory and inhibitory neurons

Models of gamma oscillations in cortical circuits generally fall into two classes. One that generates oscillations with a single pool of inhibitory cells, and one that generates oscillations by the reciprocal connections between pools of inhibitory and excitatory principal neurons (Tiesinga and Sejnowski, 2009; Whittington et al., 2000). While it is generally thought that the former class captures gamma oscillations in the dentate gyrus (Vida et al., 2006; Espinoza et al., 2018; Ewell and Jones, 2010; Diamantaki et al., 2014), we wanted to test whether our findings also generalize to networks of coupled excitatory and inhibitory neurons.

To test this, we built a model composed of 200 anatomically detailed PV neurons and 800 principal neurons that were synaptically coupled based on empirical data (Fig. 6a, cartoon, see Methods Ring networks of PV basket cells and principal cells). Both neuronal populations received Poisson type synaptic stimulation. We considered again two cases that depended on whether PV neurons were driven by input close to the soma (Fig. 6a-c) or on the dendrites (Fig. 6d-f). In networks with homogeneous input across neurons, synchrony among PV cells and, to a lesser degree, principal neurons emerged rapidly, but as the input heterogeneity increased, synchrony was only maintained when PV cells received input on the dendrites (compare Fig. 6c and f). Moreover, even in homogeneous networks (0% input heterogeneity cases) dendrite-driven networks showed a higher synchrony. This further illustrates, as discussed earlier, that PV neuron dendrites not only buffer spatial input heterogeneities but also temporal heterogeneities in a variety of network architectures, leading to high neuronal synchrony.

**Figure 6.**
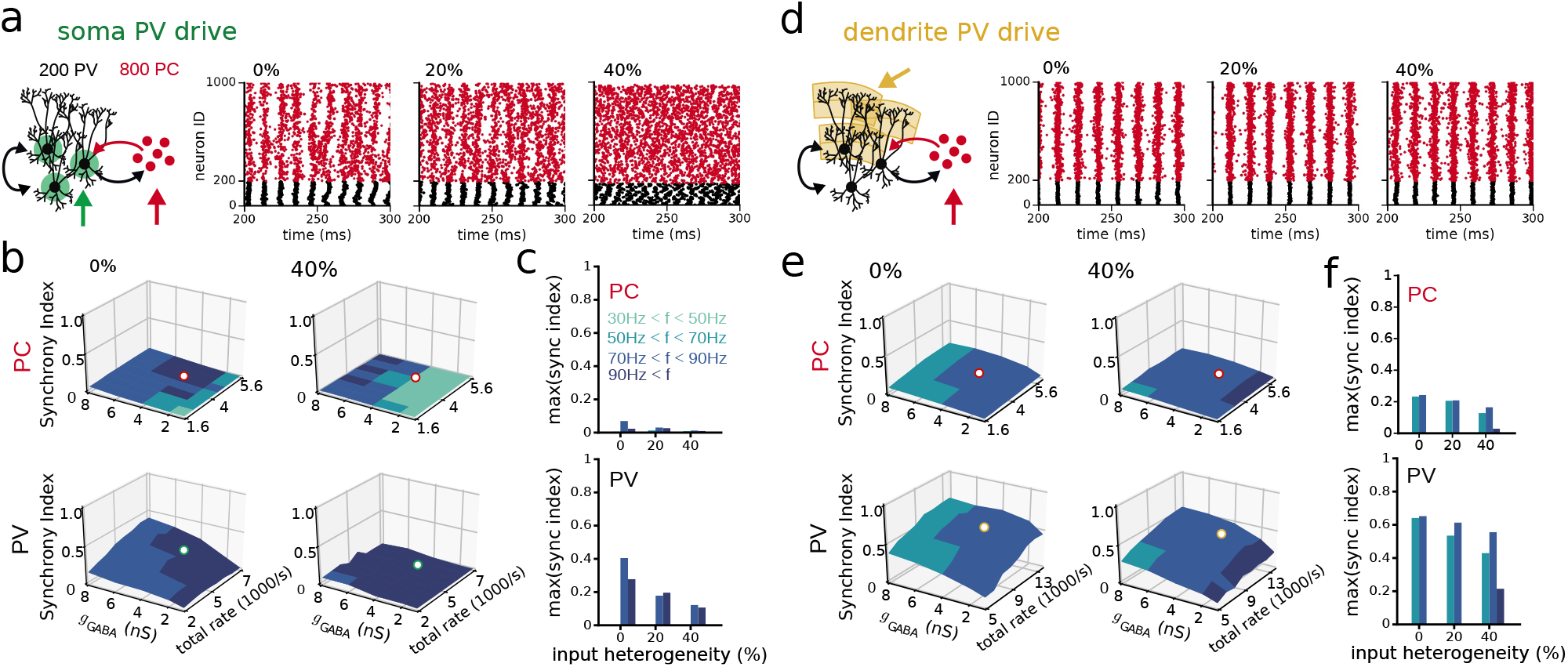
PV dendrites enhance synchrony in networks of synaptically coupled excitatory and inhibitory neurons. **(a)** Simulations using networks of reciprocally coupled PV neurons and principal cells (PC, see Methods Ring networks of PV basket cells and principal cells). The PV neurons are also coupled among themselves. Both populations of cells are driven by an excitatory drive with varying mean rate and input heterogeneity (green and red arrow). The external drive to PV cells is located perisomatically (green shaded area). Spike raster diagrams show activity of 200 PV cells (black) and 800 PCs (red) for 0%, 20% and 40% input heterogenety. **(b)** The Synchrony Index as a function of inhibitory coupling strength (between PV cells) and total input rate to PV cells for 0% and 40% input heterogeneity (top row: PCs, bottom row: PV cells). The dots on the Synchrony Index corresponds to the examples in (a). **(c)** The maximum Synchrony Index per frequency band, as a function of input heterogeneity (PCs, top; PV cells, bottom). **(d-f)** Same as (a-c), but now using dendritic input to PV cells (yellow shaded area).

## Discussion

We show that PV neuron dendrites are critically important for enhancing synchronous activity at gamma frequencies in heterogeneous networks. We found that the dendrites scale down the gain of the I-O relationship and reduce the cell’s sensitivity to input fluctuations due to the high levels of dendritic K^+^-channels. Anatomically detailed network models reveal that these properties help to homogenize firing rates so that PV neurons can synchronize at a common frequency. Therefore, we propose that the unique biophysical properties of PV neuron dendrites enhance the robustness of gamma oscillations.

For decades, experiments and theory tried to explain how spike synchrony emerges in heterogeneous networks. Classic work studying gamma oscillations in vitro, by perfusing brain slices with excitatory receptor agonists (Whittington et al., 1995; Fisahn et al., 1998), showed that fastspiking cells can synchronize when their input varies between ~ 35% to ~ 53% (Vida et al., 2006). However, this likely underestimates the conditions in vivo when synaptic input is more cell-selective and noisier (Destexhe et al., 2003), favoring desynchronization. Early network models tolerated only 3-5% heterogeneity of the tonic drive (Wang and Buzsáki, 1996), until reports showed that inhibitory connections between PV neurons are faster and stronger than previously thought (Bartos et al., 2002, 2001). Based on these data, network models could tolerate a heterogeneous tonic drive up to 10% (Bartos et al., 2002). A landmark study showed that shunting inhibition further enhances robustness and increases tolerance to heterogeneous input of 30-70% (Vida et al., 2006). However, the enhancing effects of shunting inhibition work only under restrictive conditions (Kotani et al., 2014; Tikidji-Hamburyan and Canavier, 2020a), and simulations show that shunting inhibition depends on low excitatory drive and strong inhibitory coupling (Vida et al., 2006) (Suppl. Fig. S2a-c). Therefore, a key mechanism for synchrony in heterogeneous networks with realistic inhibitory coupling was lacking.

By considering the dendrites, we show that networks can tolerate high levels of input heterogeneity, well beyond 100%. This finding is independent of whether inhibition is hyperpolarizing or shunting and relaxes the requirement for strong inhibitory coupling by an order of magnitude (Suppl. Fig. S2c). We also increased biological realism to the model network by adding other forms of heterogeneity. Previous models mainly used tonic excitation. Instead, we used Poisson trains of synaptic conductances, adding a significant amount of synaptic noise that reduces synchrony (Suppl. Fig. S3d). We also used networks composed of different cell models, each with a unique morphology and excitability, resembling biological networks (Suppl. Fig.S5f). Finally, we show that PV cells in highly heterogeneous networks can synchronize during hippocampal theta oscillations (Fig. 5) and in different network architectures (Fig. 6). In summary, PV dendrites strongly enhance spike synchrony in inhibitory networks under a wide range of conditions, which is necessary to withstand the plethora of heterogeneities that is typical for biological networks.

What are the biophysical mechanisms that enable PV neuron dendrites to enhance spike synchrony? We find that two factors play a role. First, PV neuron dendrites lack regenerative events such as Na^+^ and Ca^2+^-spikes (Hu et al., 2010), and have only low levels of NMDA receptors (Koh et al., 1995b). Instead, they are dominated by high-threshold and fast-activating K^+^-conductances (Hu et al., 2010). Second, thin PV dendrites have a high input impedance and rapidly depolarize when driven by synaptic input. This reduces the driving force for excitation and limits the amount of synaptic current that can be generated (Bush and Sejnowski, 1994). While both factors reduce the gain of the I-O relationship and the amplitude of membrane potential fluctuations, the dendritic K^+^-conductances play a more important role (Fig. 4). Yet, surprisingly, K^+^-channels are not necessary. Without K^+^-channels, PV neuron networks remain robust to heterogeneities because of the enhanced contribution of a reduction in driving force (Fig.4g). These properties of PV neurons in the dentate gyrus may not be exceptional. There is evidence that the dendrites of PV cells in other cortical circuits have similar properties (reviewed in Hu and Vervaeke (2018), but see Chiovini et al. (2014)). Furthermore, rhythm-generating inhibitory neurons in the cerebellar cortex, such as stellate (Abrahamsson et al., 2012) and Golgi cells (Vervaeke et al., 2012) have also thin, high impedance dendrites lacking regenerative properties, and the sub-linear integration in these cells is dominated by a reduction in synaptic driving force. Therefore, we speculate that some inhibitory interneurons have dendrites that are ideally suited for rhythm generation. Altogether, this illustrates that care should be taken when simplifying inhibitory neurons as point neurons without dendrites in network models (Tzilivaki et al., 2019; Poirazi and Papoutsi, 2020).

Neurons integrate input from different origins that is spatially segregated on their dendrites. A corollary of our results is that input from the entorhinal cortices on the outer two-thirds of the apical dendrites is more likely to generate gamma oscillations compared to the commissural input that targets the proximal dendrites. Spatial segregation of synaptic inputs on the dendrites is common in the brain. For example, PV cells in the layer 4 of the neocortex receive thalamic inputs close to the soma, while intra-cortical contacts are rather located on the distal dendrites (Freund et al., 1985; Bagnall et al., 2011). Depending on whether the cortical circuit is dominated by sensory or intracortical activity, this may promote network desynchronization and synchronization, respectively. In conclusion, our results suggest that the biophysical properties of PV neuron dendrites promote spike synchrony in the gamma frequency range, and support the many cognitive functions associated with this rhythm.

## ACKNOWLEDGEMENTS

We thank the members of the Vervaeke lab for discussions and comments on the manuscript, and Padraig Gleesson for his continuous support of neuroML. This work was supported by an ERC starting grant (639272) to KV, a Young Research Talents grants from the Norwegian Research Council to KV (513401), a Toppforsk grant from the Norwegian Research Council (#276047), an EEA grant (RO-NO-2019-0504), and EU Seventh Framework Scientia Fellows fellowship to BK ((609020, FP-PEOPLE-2013-COFUND). HH was supported by the Norwegian Research Council (#250866). We thank Aaron Millstein, Raul Muresan, Christoph Schmidt-Hieber and Alexandra Tran-Van-Minh for providing critical comments that significantly improved a draft of the manuscript.

## AUTHOR CONTRIBUTIONS

BK and KV designed the study. BK performed all simulations and analyses. HH performed all experiments. BK and KV wrote the paper. All authors discussed the results.

## Methods

### Experimental procedures

#### Dendritic patch-clamp recordings

Experiments were ethically approved by the Norwegian Food Safety Authority (Mattilsynet), and were performed in strict accordance with institutional, national, and European guidelines for animal experimentation.

Transverse hippocampal slices (thickness 350 μm) were prepared from brains of 17- to 23-day-old male Wistar rats. Rats were housed under a 12h light (7am-7pm) and dark (7pm-7am) cycle and were kept in a litter of 8-10 animals together with the mother in a single cage. Slices were cut in ice-cold, sucrose-containing physiological extracellular solution using a vibratome (VT1200, Leica Microsystems), incubated in a storage chamber filled with standard physiological extracellular solution at ~ 34°C for 30 min, and subsequently stored at room temperature. Slices were then individually transferred into a recording chamber and perfused with standard physiological extracellular solution containing 125 mM NaCl, 25 mM NaHCO_3_, 2.5mM KCl, 1.25mM NaH_2_PO_4_, 2mM CaCl_2_, 1 mM MgCl_2_, and 25 mM D-glucose (equilibrated with 95% O_2_ and 5% CO_2_ gas mixture). Current-clamp recordings were performed at near-physiological temperature (~ 33^°^C; range: 31-34^°^C).

For recordings from interneuron dendrites, we used the following strategy. First, a somatic recording was obtained using an internal solution containing Alexa Fluor 488 (50 or 100 μM, Invitrogen). Second, after ~ 30 min of somatic whole-cell recording, the fluorescently labeled axon and dendrites were traced from the PV basket cell soma with a Nipkow spinning disk confocal microscope (Volocity, Perkin Elmer, equipped with an Orca camera, Hamamatsu and a solid-state laser with excitation wavelength of 488 nm). Total exposure time was minimized to avoid photodamage. Finally, fluorescent and infrared differential interference contrast (IR-DIC) images were compared and dendrites were patched under IR-DIC.

Patch pipettes were fabricated from thick-walled borosilicate glass capillaries (outer diameter: 2 mm, inner diameter: 1 mm) with a horizontal pipette puller (P-97, Sutter Instruments). The intracellular solution for soma-dendritic whole-cell recordings contained 120 mM K-gluconate, 20mM KCl, 10mM EGTA, 2mM MgCl_2_, 2mM Na2ATP and 10mM HEPES, pH adjusted to 7.3 with KOH. When filled with internal solution, they had a resistance of 2-10 MΩ for somatic recordings and 6-40 MΩ for dendritic recordings. Current-clamp recordings were performed using a Multiclamp 700B amplifier (Molecular Devices). Series resistance was 12-90 MΩ. Cells with somatic resting potentials more positive than −50 mV were discarded. Pipette capacitance and series resistance compensation (bridge balance) were applied throughout experiments. The input-output relationship was determined by injecting 1-second depolarizing current pulses of various amplitudes into the soma at the resting membrane potential (−60.7± 0.8 mV). The resistance of patch pipttes was between 6.7 and 11.6 MΩ.

Signals were sampled at 50 or 100 kHz with a Digidata 1322 converter board (Molecular Devices) and low-pass filtered at 10 kHz. Data acquisition and generation of pulse protocols were performed with pClamp 9 or 10 (Molecular Devices).

PV basket cells were identified based on the nonaccommodating, fast-spiking phenotype (steady-state spike frequency >150 Hz at physiological temperature in response to 1-s, 0.3- to 1-nA somatic current pulses), and the morphological properties of the axonal arbor, which was largely restricted to the granule cell layer and established basket-like structures around granule cell somata that were visible in the confocal images. In a previous publication (Hu et al., 2010) a large sample of fast-spiking interneurons in the dentate gyrus was analyzed in detail by light microscopy. It was concluded that the fast-spiking phenotype was tightly correlated with the expression of parvalbumin (Hu et al., 2010). Furthermore, 78 of 83 cells were classical basket cells with tangential axon collaterals and basket-like branches around granule cell somata, whereas 5 out of 83 were axo-axonic cells with radial axon collaterals (Hu et al., 2010). Based on these results, the recorded cells were termed PV basket cells throughout the present study.

To describe the propagation of EPSPs from PV basket cell dendrites to the soma (Fig. 2a), miniature EPSPs were evoked by injecting a high osmotic external solution to the proximity of the dendritic recording site during simultaneous soma–dendrite recordings with a third patch pipette. 30 μM ZD7288 and 2 μM SR95531 were added to the standard physiological solution in these experiments to block HCN channels and GABAA receptors. The high osmotic external solution contained 125 mM NaCl, 25 mM NaHCO3, 2.5 mM KCl, 1.25 mM NaH2PO4, 2mM CaCl2, 1 mM MgCl2, 25mM D-glucose, 300mM sucrose, 50 μM DL-APV, 30 μM ZD7288, 2 μM SR95531 and 1 μM tetrodotoxin.”

### PV Basket Cell Model

#### Passive and active properties of PV basket cell models

##### Passive parameters

A previous study (Nörenberg et al., 2010) meticulously constrained the passive parameters of the five PV basket cell models based on experimental data, see Suppl. Fig. S1a and https://senselab.med.yale.edu/ModelDB/showmodel.cshtml?model=140789. We summarize the membrane capacitance *C*_m_, axial resistance *R*_i_ leak reversal potential *E*_leak_ and membrane conductances *g*_leak_ for each of the models in Table 1 and 2. The leak conductance *g*_leak_ is spatially inhomogeneous along the dendrites (higher for closer distance *d* to the soma, lower more distally), as this was found to fit the passive response properties of dentate gyrus PV cells best (Nörenberg et al., 2010). In particular,

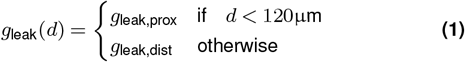

**Table 1.** Passive parameters of the detailed PV cell models. Adapted from Nörenberg et al. (2010).

| Passive parameters |  |  |  |  |  |
| --- | --- | --- | --- | --- | --- |
| Cell | PV1 | PV2 | PV3 | PV4 | PV5 |
| $C_m$ [$\mu\text{F}/\text{cm}^2$] | 0.99 | 1.06 | 0.77 | 0.92 | 0.97 |
| $R_i$ [$\Omega\text{cm}$] | 149 | 137 | 141 | 178 | 258 |
| $E_{\text{leak}}$ [mV] | -80 | -73 | -80 | -75 | -75 |

**Table 2.**
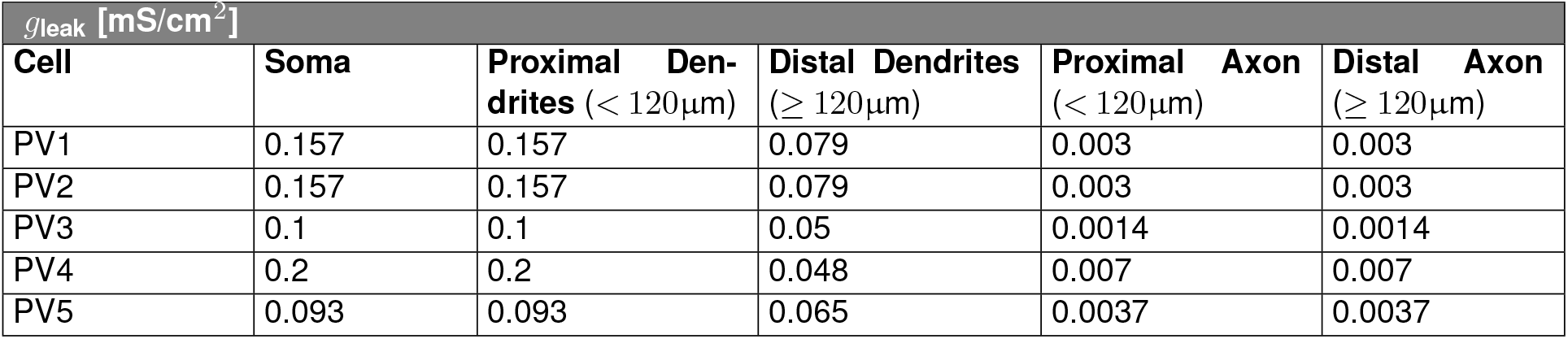
Leak conductances of the detailed PV cell models. Adapted from Hu et al. (2010); Nörenberg et al. (2010).

| $g_{\text{leak}}$ [$\text{mS}/\text{cm}^2$] | | | | | |
| --- | --- | --- | --- | --- | --- |
| Cell | Soma | Proximal Dendrites ($< 120\mu\text{m}$) | Distal Dendrites ($\geq 120\mu\text{m}$) | Proximal Axon ($< 120\mu\text{m}$) | Distal Axon ($\geq 120\mu\text{m}$) |
| PV1 | 0.157 | 0.157 | 0.079 | 0.003 | 0.003 |
| PV2 | 0.157 | 0.157 | 0.079 | 0.003 | 0.003 |
| PV3 | 0.1 | 0.1 | 0.05 | 0.0014 | 0.0014 |
| PV4 | 0.2 | 0.2 | 0.048 | 0.007 | 0.007 |
| PV5 | 0.093 | 0.093 | 0.065 | 0.0037 | 0.0037 |

##### Voltage-dependent currents

The voltage-dependent currents *I*_Na_, *I*_K1_ and *I*_K2_ are Hodgkin-Huxley-type and based on the Wang-Buzsaki model (Wang and Buzsáki, 1996; Hu et al., 2010). Here, *I*_K1_ has the same parameterization as in (Wang and Buzsáki, 1996; Hu et al., 2010), while *I*_K2_ is modified from (Wang and Buzsáki, 1996; Hu et al., 2010) to match the more depolarized activation threshold in the distal dendrites (see Fig. 1e).

###### Sodium current I_Na_

The sodium current *I*_Na_ has an activation variable *m* and an inactivation variable *h*. The dynamics of *m* is assumed to be fast and substituted by the steady-state value *m*_∞_ (Wang and Buzsáki, 1996). In particular,

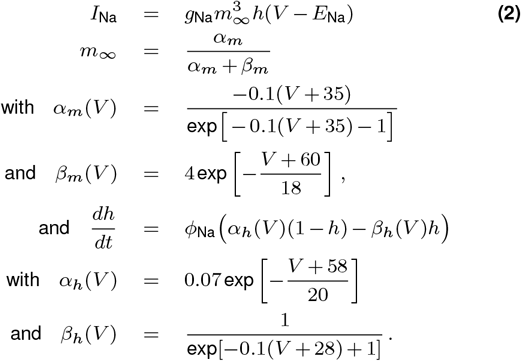

###### Potassium current I_K1_

The potassium current *I*_K1_ has activation variable *n*_1_, such that

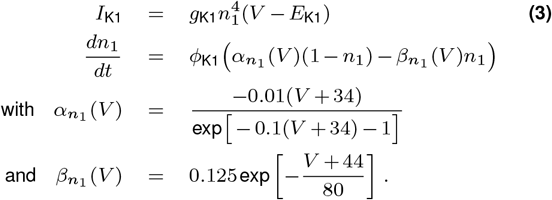

###### Potassium current I_K2_

The potassium current *I*_K2_ has activation variable *n*_2_, such that

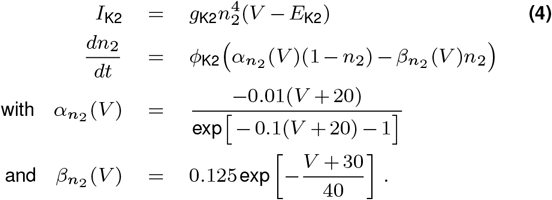

Parameters *φ, E* and *g* are summarized in Table 3 and 4.

**Table 3.** Parameters of voltage-dependent channels.

| Active Channel Parameters |  |
| --- | --- |
| $E_{Na}$ [mV] | 55 |
| $\phi_{Na}$ | 5 |
| $E_{K1}$ [mV] | -90 |
| $\phi_{K1}$ | 2 |
| $E_{K2}$ [mV] | -90 |
| $\phi_{K2}$ | 5 |

**Table 4.** Fitted parameters of voltage-dependent channels of the detailed PV cell models. Channel density of sodium *g*_Na_ and the two potassium channels *g*_K1/2_.

| Active Channel Densities |  |  |  |  |  |
| --- | --- | --- | --- | --- | --- |
| Densities [mS/cm<sup>2</sup>] | Soma | Proximal Dendrites | Distal Dendrites | Proximal Axon | Distal Axon |
| **PV1** |  |  |  |  |  |
| $g_{Na}$ | 100 | 5 | 5 | 400 | 35 (>120μm) |
| $g_{K1}$ | 40 | 40 | 0 (>200μm) | 40 | 40 |
| $g_{K2}$ | 0 | 0 | 30 (>200μm) | 0 | 0 |
| **PV2** |  |  |  |  |  |
| $g_{Na}$ | 250 | 5 | 5 | 550 | 35 (>120μm) |
| $g_{K1}$ | 50 | 50 | 0 (>200μm) | 50 | 50 |
| $g_{K2}$ | 0 | 0 | 30 (>200μm) | 0 | 0 |
| **PV3** |  |  |  |  |  |
| $g_{Na}$ | 200 | 5 | 5 | 400 | 35 (>120μm) |
| $g_{K1}$ | 50 | 50 | 0 (>200μm) | 50 | 50 |
| $g_{K2}$ | 0 | 0 | 30 (>200μm) | 0 | 0 |
| **PV4** |  |  |  |  |  |
| $g_{Na}$ | 120 | 5 | 5 | 400 | 35 (>120μm) |
| $g_{K1}$ | 60 | 60 | 0 (>200μm) | 60 | 60 |
| $g_{K2}$ | 0 | 0 | 30 (>200μm) | 0 | 0 |
| **PV5** |  |  |  |  |  |
| $g_{Na}$ | 150 | 5 | 5 | 450 | 35 (>120μm) |
| $g_{K1}$ | 50 | 50 | 0 (>200μm) | 50 | 50 |
| $g_{K2}$ | 0 | 0 | 30 (>200μm) | 0 | 0 |

Activation curves 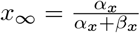 and time constants 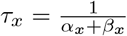 of the variables *x* = {*m, h, n*_1_, *n*_2_} as function of *V* are plotted in Fig. M1.

**Figure M1.**
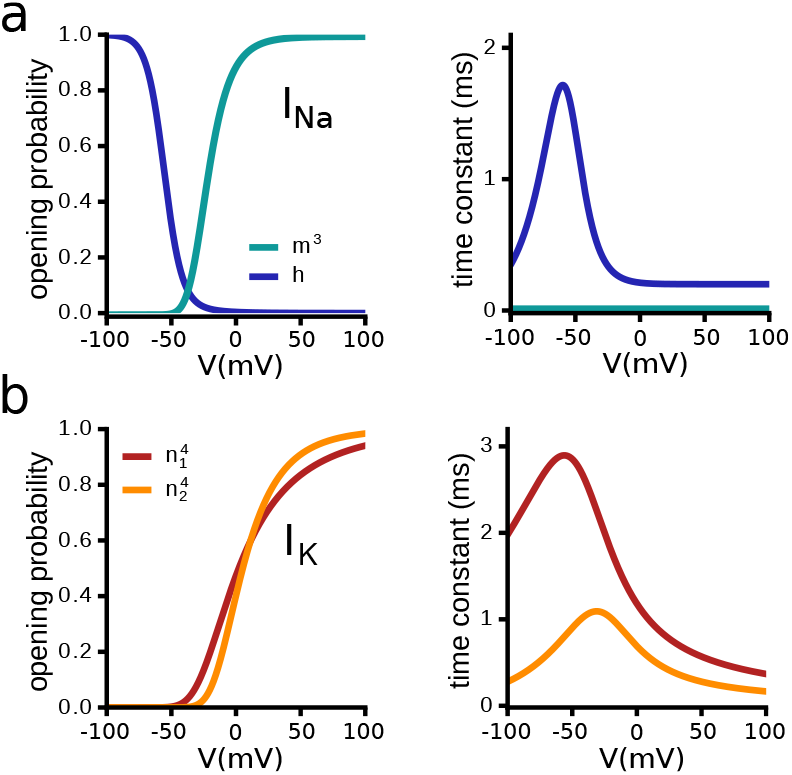
Voltage-dependent steady-state activation curves and time constants of *I*_Na_ and *I*_K1/2_. Left side: Steady state activation of the **(a)** activation variable *m*^3^ and inactivation variable *h* of the sodium channel, and **(b)** of the activation variables *n*_1_, *n*_2_ of the potassium channels. Right side: respective time constants.

###### Model fitting

For all five reconstructed basket cell models, we used the passive properties as measured in Nörenberg et al. (2010), see Tables 1, 2. Next, we systematically varied *g*_Na_, *g*_K1_ and *g*_K2_ to find good fits to the average I-O relationship that was experimentally measured using somatic current injections (Fig. 1c). Next, we conducted simulations to test whether the model matches several other experimental observations, in particular the dendritic I-O relationship (Fig. 1d), the steady-state membrane potential responses to dendritic (Fig. 1e) and somatic current injections, the spike attenuation along the apical dendrites (not shown), and the attenuation of EPSPs (Fig.2b). If necessary, we slightly adapted *g*_Na_ and *g*_K1_ to obtain a better fit to the experimental data. The resulting channel densities are listed in Table 4. Compared to *I*_K1_, we shifted the steady-state activation curve of *I*_K2_ to more depolarized potentials to obtain a better fit to the sublinear membrane potential in response to current injections in the dendrites (Fig. 1e). Note that using only *I*_K1_ on the entire dendritic tree has no significant effects on any of the main results apart from a worse fit of the dendritic V-I-curve. After introduction of the voltage-dependent channels the input resistance of the models remained relatively close to the experimental observation in Nörenberg et al. (2010), see Table 5.

**Table 5.** Input resistances: experiment vs.model. Input resistance as measured in cell models in comparison to experimental data (Nörenberg et al., 2010).

| Input Resistance |  |  |  |  |  |
| --- | --- | --- | --- | --- | --- |
| Cell | PV1 | PV2 | PV3 | PV4 | PV5 |
| $R_{inp}$ [MΩ] exp (Nörenberg et al., 2010) | 119.2 | 57.3 | 118.8 | 63.4 | 60.7 |
| $R_{inp}$ [MΩ] model | 97.5 | 63.1 | 108.3 | 48.7 | 64.5 |

### Ball-and-sticks model

This model comprises one soma compartment (diameter=length=25 μm) and five equal dendritic compartments (diameter=1 μm, length=300μm). Passive properties are *C*_m_=1μF/cm^2^, *g*_leak, soma_=0.16 mS/cm^2^, *g*_leak, dend_=0.08 mS/cm^2^, *E*_leak_=−75mV, and *R*_i_=100 Ωcm. The soma has *I*_K_ (*g*_K_=20 mS/cm^2^, *E*_K_=−90mV, *ϕ*_K1_=5 in Eqn. (3)) and *I*_Na_ (*g*_Na_=80 mS/cm^2^, *E*_Na_=55mV, *ϕ*_Na_=5 in Eqn. (2)). The dendrites have *I*_K_ (*g*_K_=20 mS/cm^2^, *E*_K_=−90mV, *ϕ*_K1_=5 in Eqn. (3)). For comparison to experimental data, see Suppl. Fig.S6a.

### Principal cell model

The regular-spiking principal cell model was adapted from Strüber et al. (2017) and can be downloaded on ModelDB (https://senselab.med.yale.edu/ModelDB/showmodel.cshtml?model=229750).

In short, it comprises one soma compartment (diameter=length=5.642 μm) with passive parameters *C*m=1.01 μF/cm^2^, *g*_leak_=0.1 mS/cm^2^, *E*_leak_=−75mV, and *R*_i_=194Ωcm. Active properties comprise a voltage-gated Na^+^ (*g*_Na_=10 mS/cm^2^, *E*_K_=55 mV), and a delayed-rectifier voltage-gated K^+^-conductance (*g*_K_=15 mS/cm^2^, *E*_K_=−90 mV; the original model in Strüber et al. (2017) had three K^+^-conductances, delayed rectifier, A-type, and M-type, however, only the delayed rectifier had non-zero peak-conductance in the published code on ModelDB, so we adopted this choice).

### Network models

#### Ring networks of PV basket cells

The ring networks are based on previous models of gamma oscillations (Bartos et al., 2002; Vida et al., 2006) (for overview of network connectivity, see Appendix Simulation parameter tables, Table 7). We arranged 200 PV basket cells along a ring where the distance between neighboring cells is 50 μm. The probability that two neurons are coupled by inhibitory synapses follows a Gaussian distribution *p*_ring_ (*d*) = *e*^−*d*2/2σ2^, with *d* being the distance between the cell somata, and σ= 1 200 μm (Bartos et al., 2002; Vida et al., 2006) (Fig. M2a). Synaptic connections are not allowed between cells that are more than 2 500 μm apart. According to these connectivity rules, each PV basket cell is connected to approximately 58 other PV basket cells (Bartos et al., 2002; Vida et al., 2006) (see Appendix Ring networks for a derivation). These rules are constrained by the PV basket cell density and the extent of their axonal tree in hippocampal area CA1 (Sik et al., 1995).

When two cells are synaptically connected, we randomly distributed *n* synapses in the perisomatic region (≤ 50 μm from soma). The number of synapses between two PV basket cells was distance-dependent following the same equation as the coupling probability, but with a maximum of five synapses. The number of synaptic contacts was rounded down to an integer (Fig. M2b, more details in Appendix Ring networks).

**Figure M2.**
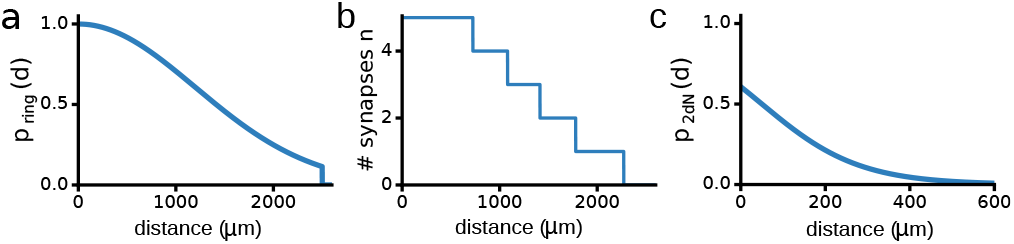
Connection probabilities. **(a)** Connection probability in the ring networks (Bartos et al., 2002; Vida et al., 2006). **(b)** Number of synaptic contacts as a function of distance. **(c)** Connection probability in the two-dimensional network. Curve adapted from Espinoza et al. (2018).

##### GABAergic synaptic conductances

We modeled synaptic conductances as two-state kinetic scheme synapses (“Exp2Syn” model in NEURON) i.e.,

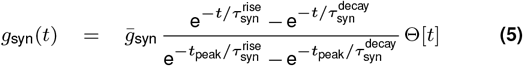

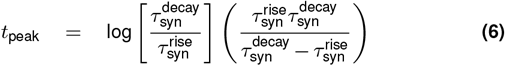

For GABAergic inhibitory synapses, the rise time constant 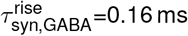 (Bartos et al., 2002; Vida et al., 2006), and the decay time constant 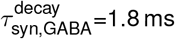 (Bartos et al., 2002; Vida et al., 2006) (for overview of parameters, see Appendix Simulation parameter tables, Table 6). We varied the peak amplitude of the individual GABAergic synaptic conductances 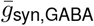 between 0.5 and 10 nS (≈ 0.008-0.08 mS/cm^2^), with a synaptic reversal potential *E*_syn,GABA_=−75 mV for hyperpolarizing inhibition, and −60 mV for shunting inhibition (for shunting inhibition, we varied 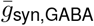 up to 30 nS, because synchronous states typically require very strong inhibition (Vida et al., 2006; Tikidji-Hamburyan and Canavier, 2020b)). Synaptic currents are described by IPSC(*V, t*) = *g*_syn,GABA_(*t*)(*V* — *E*_syn,GABA_).

**Table 6.** Parameters of “Exp2Syn”-synapses (Eqn. (5)).

| Synaptic Parameters |  |  |  |  |
| --- | --- | --- | --- | --- |
| Synapse Type | $\tau_{\text{syn}}^{\text{rise}}$ [ms] | $\tau_{\text{syn}}^{\text{decay}}$ [ms] | $\bar{g}$ [nS] | $E_{\text{rev}}$ [mV] |
| GABA (PV→PV) | 0.18 | 1.8 | 0.5-10 | −75 (hyperpol), −60 (shunting) |
| AMPA | 0.2 | 2 | 2 | 0 |
| NMDA | 3 | 70 | 0.44 | 0 |
| PV→PC | 0.16 | 7 | 0.1 | −65 |
| PC→PV | 0.1 | 1 | 2 | 0 |

The spike detection threshold of PV basket cells and the Ball-and-Stick model was set to 0mV. The delay *τ* of spike transmission comprised a constant synaptic delay of *τ*_0_=0.5ms (Bartos et al., 2002; Vida et al., 2006) and a distance dependent conduction delay of *τ*(*i*)=0.2·*i* ms (Bartos et al., 2002; Vida et al., 2006), where *i* is the absolute difference between the indices of two neurons on the ring network. Given that neighboring neurons are separated by 50 μm, this corresponds to a conduction velocity of 0.25 m/s (Bartos et al., 2002; Strüber et al., 2017). In a network of 200 cells, the conduction delays varied between between 0.7 ms and 10.5 ms, with an average delay 〈*τ*〉 ≈ 4.1 ms (see also Appendix Ring networks). To avoid onset transients, the network synapses were only activated after 150 ms.

##### Electrical synapses

Where mentioned, we also included electrical synapses. We positioned ten gap junctions of 0.1 nS (Bartos et al., 2002; Vida et al., 2006) randomly on the perisomatic area (≤ 50μm from soma), and coupled each cell to four out of eight randomly chosen nearest neighbors (four to the left, four to the right). Therefore, gap junctions were only present between cells that were at most 200 μm apart (Espinoza et al., 2018). We tested the strength of gap junction coupling by measuring the coupling coefficient between electrically coupled cells which was typically around 9% (Hormuzdi et al., 2001; Venance et al., 2000) (Suppl. Fig. S4a).

#### Ring networks of PV basket cells and principal cells

Networks of coupled excitatory and inhibitory cells consisted of *N*_I_=200 PV basket cells and *N*_E_=4*N*_I_ principal cells (PC) arranged on two rings, respectively. The spatial footprint of PV→PC was the same as for PV→PV, i.e., *p*_ring_(*d*) = *ce*^*–d2/2σ2*^ with σ=1 200μm. For PV→PV connectivity, *c*=1, resulting in ~ 58 connections per cell, while for PV→PC, *c*=0.35, resulting in ~ 20 connections from PV cells to any PC (Strüber et al., 2017). The footprint from PC→PV was Gaussian with a footprint of σ=500μm and *c*=0.5, resulting in ~ 50 connections from PCs to any PV cell (Strüber et al., 2017). Synapses from PV→PV were the same as for the PV network with varied peak conductance 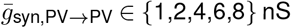. Synapses from PV→PC were “Exp2Syn”-synapses Eqn. (5) with 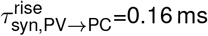 (Strüber et al., 2017), decay time constant 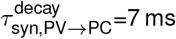 (Strüber et al., 2017), and peak conductance 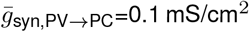, corresponding to ~ 0.5 nS for a typical dentate gyrus granule cell of size ~ 500μm^2^ (Clairborne et al., 1990). Synapses from PC→PV were AMPA-type “Exp2Syn”-synapses with 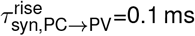 (Strüber et al., 2017), decay time constant 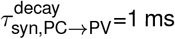 (Strüber et al., 2017), and peak conductance 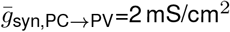 (Strüber et al., 2017) (for overview of parameters, see Appendix Simulation parameter tables, Table 6). Spike detection threshold was set to 0 mV for PV basket cells, and to −20 mV for principal cells.

#### Two-dimensional PV basket cell network

In addition to ring networks, we also created two-dimensional networks based on recent empirical data describing the distance dependence of the coupling probability between PV basket cells in the dentate gyrus (Espinoza et al., 2018) (for overview of network connectivity, see Appendix Simulation parameter tables, Table 7). It is given by the following equation and plotted in Fig. M2c:

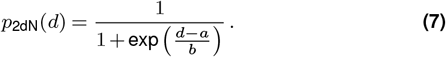

**Table 7.** Specifications of network models.

| Network Specifications |  |  |  |  |  |
| --- | --- | --- | --- | --- | --- |
| Network Type | Dimension | Metric | Boundary Condition | Cell Distribution | Connectivity Profile |
| ring network | one | absolute difference | periodic | equilateral grid | boxcar<br>$p(d) = \Theta[d_{\text{max}} - d]$ |
| random network | two | euclidean | periodic | uniform random | sigmoid<br>$p(d) = \frac{1}{1 + \exp(\frac{d-a}{b})}$ |

Here, we used *a*=50 and *b*=115 to approximately match the connection probability fit in Espinoza et al. (2018). Notably, the spatial reach of connectivity is much smaller compared to the ring networks (Fig. M2a,c), which means that assuming the same conduction velocity as used in ring networks, i.e., 0.25 m/s, the effective delays will be much shorter. Therefore, the effect of synchronized inhibition will be shortened and oscillations will be faster. To compare synchrony in both network types in terms of network structure only, we thus assumed a three-times slower conduction velocity (0.0833 m/s) to match compound inhibition *G*_GABA_(*t*) in fully synchronous networks (for a detailed derivation see Appendix Ring networks). Neurons were distributed uniform-randomly on a torus to avoid boundary effects.

#### External drive

##### PV ring and 2d-networks

In most simulations, neurons were driven by Poisson-type excitatory synapses, that were distributed uniformly either perisomatically (<50μm from the centre of the soma) or on the outer two-thirds of the apical dendrites, i.e., ≳120μm (≳150μm for ball-and-stick cells) from the soma centre. The total input rate *r*_stim_ = *n*_syn_ × *r*, where *n*_syn_ is the total number of synapses and *r* is the rate per synapse. We did not observe any qualitative or quantitative differences when varying *n*_syn_ or *r* while keeping *r*_stim_ fixed. Therefore, we decided to use *n*_syn_=50 for perisomatic, and *n*_syn_=100 for distal apical drive, and varied *r* between 20 and 200 Hz.

In most cases the synapses were AMPA-type synapses modeled as “Exp2Syn”-synapses, see Eqn. (5). In the model, we used a rise time constant 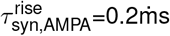 and a decay time constant of 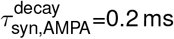, consistent with previous data (Geiger et al., 1997; I.C. and H.P., 1999), and our own measurements in Fig. 2. The peak conductance was 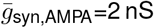 (except in the ball and stick model, where it was 0.5 nS). The synaptic reversal potential was 0 mV. The postsynaptic currents are given by EPSC((*V, t*) = *g*_syn,AMPA_(*t*)(*V* — *E*_syn,AMPA_), see overview Appendix Simulation parameter tables, Table 6.

##### PV basket cell-PC networks

PV basket cells and PCs were driven by Poisson-type excitatory AMPA-type synapses (“Exp2Syn”-synapses, *τ*^rise^=0.1 ms (Strüber et al., 2017), decay time constant *τ*^decay^=1 ms (Strüber et al., 2017), peak conductance 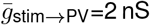, 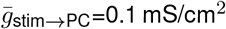, corresponding to ~ 0.5 nS for a typical dentate gyrus granule cell of size ~ 500μm^2^ (Clairborne et al., 1990)). Input rates were covaried, such that input rates to PCs ranged from *r*_stim,PC_=2 000-6 000/s, and rate per synapse for PV basket cells was *r*_stim,PC_/40 (100 synapses for dendritic stimulation, 50 synapses for perisomatic stimulation).

##### AMPA/NMDA synapses

We also simulated co-localized AMPA/NMDA synapses (Fig. S4d-f). The NMDA-conductance is given by an exponential rise and decay time constant, and is voltage-dependent to model the Mg^2+^-block (Farinella et al., 2014) as (see also Eqn. (5)):

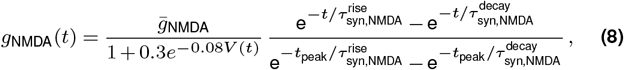

The rise and decay time constant were 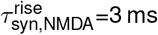 and 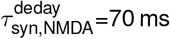 (Farinella et al., 2014), respectively. The time course of the NMDA conductance is plotted in Fig. S4d. The ratio between the maximum NMDA and AMPA conductance (0.22) and the voltage-dependence of the Mg^2+^-block (Fig. M3a,b) was based on measurements from fast spiking basket cells of the dentate gyrus reported in Koh et al. (1995a) (Appendix Simulation parameter tables, Table 6).

**Figure M3.**
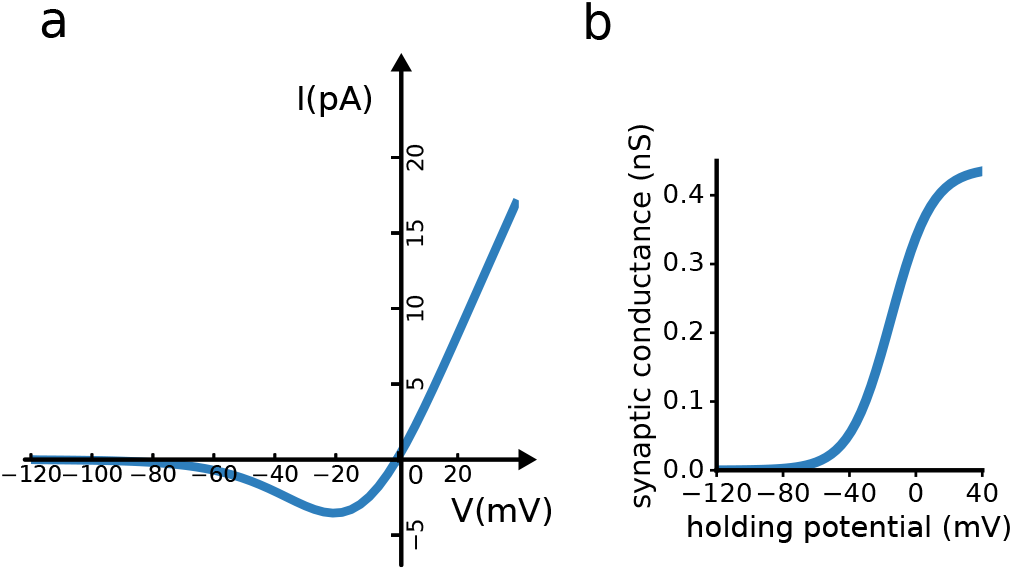
NMDA synapse. **(a)** I-V-relationship of the NMDA conductance. **(b)** Voltage dependence of the NMDA conductance.

##### Current-based synapses

Finally, to simulate distributed synaptic current sources that have the same kinetics as AMPA synapses, but that are independent of the driving force (Fig. 4c,e), we kept the membrane potential variable constant to the leak reversal potential (*E*_leak_=−75 mV for the anatomically detailed models, and −65 mV for the Ball-and-stick model). That is, EPSC(*t*) = *g*_syn,AMPA_(*t*)(*E*_leak_ — *E*_syn,AMPA_) = const × *g*_syn,AMPA_(*t*).

##### Simulations of theta-nested gamma oscillations

In order to simulate theta-modulated activity we added a sinusoidally modulated current to the somatic compartment of all cells:

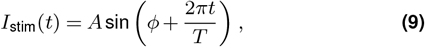

with oscillation period *f*=5 Hz, and amplitude *A*=1 nA. Additionally, all cells received a heterogeneous amount of Poisson-type excitatory synaptic input as described above.

### Simulations and analysis

#### Simulation

Single cell simulations typically covered 2 000 ms. For neurons driven with noisy or randomly distributed input we repeated the simulation ten times with different initial conditions for each parameter combination.

Network simulations typically covered 500 ms. After an initial period of 150 ms, the GABA-ergic synapses were activated. Some simulations (e.g., theta-nested gamma-oscillations) were run for 3 000 ms to obtain sufficient data. Each simulation was repeated at least 5 times with different random seeds per parameter combination to ensure different network structures, initial conditions, synapse locations and input spike trains.

The simulation time step was chosen between *δt*=0.005ms and 0.025 ms, depending on simulation requirements, e.g., to avoid numerical errors in case of rapidly fluctuating noisy inputs, or when electrical synapses were included.

All neurons and networks were simulated in NEURON/7.4 via the pyNeuroML-interface in NeuroML/v2beta4 (https://www.neuroml.org/neuromlv2). Larger network simulations were run on the Sigma2 high-performance clusters Abel and Saga (https://www.sigma2.no). Versions of the models and simulation scripts to reproduce the figures in the manuscript are available at [INSERT OpenSourceBrain-link for publication]. Data handling and analysis were done in Python/2.7.15, using the NumPy/1.11.9, SciPy/0.17.0, and Matplotlib/1.5.1 libraries.

#### Analysis

For analysis of single neuron dynamics we used the last 1 500 ms of 2 000 ms simulated time for analysis, while for the analysis of network activity we used the last 300 ms of 500 ms of the simulations (Section Simulation). For spike-based synchrony (Synchrony Index, Coherence) and regularity (CV) measures, we excluded neurons that spiked less than two times in the considered time interval. The number of neurons to average over was adapted accordingly.

##### Rate

Individual spike rates were computed as the number of spikes emitted by a neuron over the considered time interval *T*. Population spike rates were individual spike rates averaged across all neurons (spiking or not) in a network.

##### Coefficient of variation of interspike intervals

The coefficient of variation CV of interspike intervals ISI is defined as the standard deviation of its ISIs σ(ISI) over the mean of ISIs μ(ISI), i.e., CV(ISI) = σ(ISI)/μ(ISI).

##### Oscillation frequency

We computed the average membrane potential of all neurons and calculated its power spectrum. The frequency of the dominant peak then defined the oscillation frequency.

##### Synchrony Index

We first binned the spikes of all neurons that spiked at least two times in *δ*=2 ms bins to obtain the spike histrogram (Fig. M4a). Next, we calculated the Fano Factor (FF) of the spike histogram (see Fig. M4a),

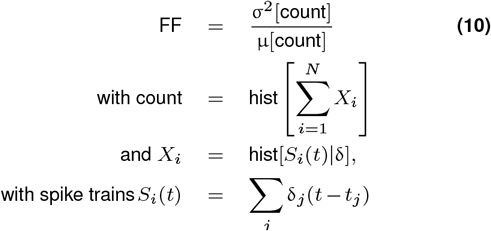

We then divided FF by the number of spiking neurons in the network. Thus, if all cells spike synchronously, the variance of the bin size will be maximal and FF/N equal to 1. If all cells spike asynchronously, the variance of the bin size will be minimal and FF/N will approach 0.

**Figure M4.**
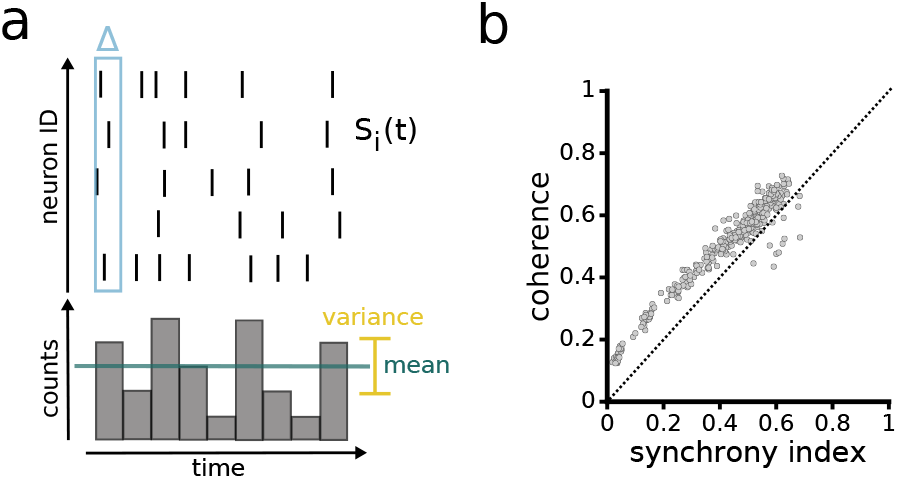
Synchrony Index: Fano Factor measures population wide synchrony. **(a)** Cartoon showing how the Synchrony Index is computed. Upper panel: spike raster plot. Lower panel: population spike count histogram. **(b)** Example scatter plot of Synchrony Index and Coherence (Bartos et al., 2002; Wang and Buzsáki, 1996).

To quantify network synchrony, previous work used the so-called Coherence Measure (Bartos et al., 2002; Wang and Buzsáki, 1996). To compare the Synchrony Index with the Coherence Measure, we plotted their relationship (Fig. M4b). Note that both measures give very similar results. However, FF/N had several advantages: First, it does not have the floor effect that we observed for coherence-based measures (Fig. M4b, see also Appendix Synchrony Index and Coherence). Second, the Synchrony Index is simpler and infers the population-wide synchrony more directly, because it is inherently a population-activity measure, whereas the Coherence Measures are based on pairwise synchrony (see Appendix Synchrony Index and Coherence for a discussion and more detailed analysis).

## Appendix

### Simulation parameter tables

Tables 6 and 7 summarize synaptic and network parameters, respectively.

### Connectivity statistics of ring and two-dimensional networks

In Section Network models we introduced the network model we generally used in the paper, the choice motivated by one of the most prominent models for studying biophysically constrained gamma-oscillations (Bartos et al., 2002; Vida et al., 2006). In some simulations (Suppl. Fig. S5a-c) however, we also simulated networks constrained by recent connectivity statistics data measured in simultaneous patch-clamp recordings of PV basket cells (Espinoza et al., 2018), see Section Twodimensional PV basket cell network. These networks differ from ring networks in several ways. First, the connection probabilities are quite different, especially with respect to the spatial reach, see Suppl. Fig. S5a,b. Second, the density of cells as a function of distance differs in onedimensional ring networks and two-dimensional torus networks. This will have pronounced impact on the number of connections at any given distance, and also on the delay distribution and, hence, timing of incoming spikes. This makes a direct comparison difficult.

In this section we will quantitatively analyze these differences in formal detail and discuss our choice to match the compound inhibitory conductance G_GABA_(*t*) each neuron would receive given total synchrony in both networks. This choice allows us to directly relate results for the two networks with an emphasis on topology.

### Ring networks

The standard network setting is derived from the networks considered in Bartos et al. (2002); Vida et al. (2006), with *N* neurons arranged equidistally on a ring. Connectivity was established randomly following a Gaussian connection probability

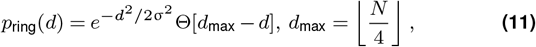

with Heaviside step-function Θ[*x*] =0 if *x* < 0, Θ[*x*] = 1 otherwise, and floor-operator ⌊*x*⌋ that rounds *x* to the next lowest integer. The standard deviation was chosen σ = 24 neuron distances (Bartos et al., 2002; Vida et al., 2006). Autapses were usually excluded, and test network runs including autapses did not show any palpable differences (not shown). In a ring network neuron density is discrete, but constant with distance, i.e.,

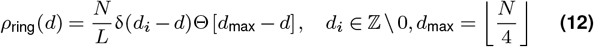

with Dirac measure *δ*(*x*) = 1 if *x* = 0, and zero otherwise. Hence the probability mass function of distances *f*(*d*) (see Fig. M5a) is given by

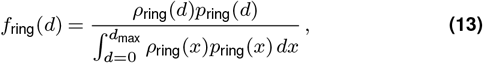

where the denominator yields the expected number of connected neurons per neuron 〈*k*〉,

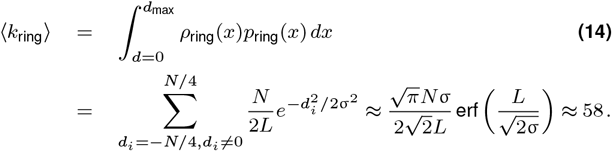

If two neurons were connected, a total number of *n*_syn_ = ⌊*n*_max_*p*(*d*)⌋ (*n*_max_ = 6, so max(*n*_syn_) = 5 in absence of autapses) GABAergic synapses were randomly distributed across a perisomatic region including the soma (with probability 1/3) and dendrites (with probability 2/3) upto 50μm from the the somatic center. This yielded an average number of synapses 〈*n*_syn_〉 ≈ 3.7 between any two connected neurons.

The distance-dependent total conductance 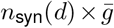 between any two neurons (also known as unitary conductance) is plotted in Fig. M5b,c for three different synaptic strengths in absolute and relative (to cell membrane area) terms, respectively.

The spike transmission delay *τ* is a linear function of the distance, i.e.,

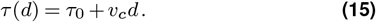

The constant offset *τ_0_* represents a synaptic delay, while the distance dependent delay is a conduction delay that represents the finite velocity *v_c_* of action potential propagation along the axon. The resulting probability mass function *f*_ring_(*τ*) of delays is given by (see Fig. M5e)

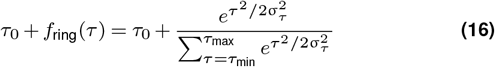

with σ_*τ*_ = 0.2σ and expectation value

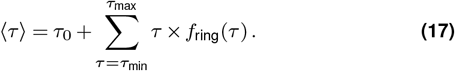

For our choice of parameters (*N*=200, *υ_c_*=0.2 m/s (Bartos et al., 2002; Strüber et al., 2017), *τ*_0_=0.5 ms) the delays in the network varied between 0.7 ms and 10.5 ms in steps of 0.2 ms, yielding an average 〈*t*〉 ≈ 4.1 ms.

If all neurons spike synchronously, and taking into account the multiplicity of synapses per connection *n*_syn_, we thus expect a compound inhibitory conductance time course (see Fig. M5d)

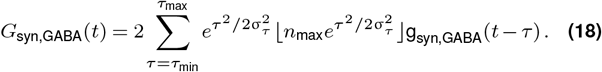

### Two-dimensional networks

To avoid boundary effects, we embedded *N* neurons in a two-dimensional torus of size *L × L*. Neuron positions were chosen uniform-randomly. Networks were created following the spatial connection probability of PV basket cells measured in Espinoza et al. (2018) (Fig. M2b, Suppl. Fig. S5a). In particular,

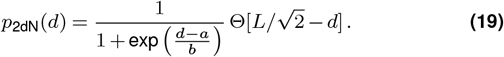

The expected number of connections per cell is then given by

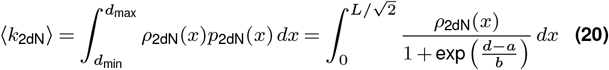

with cell density at a given distance *d* given by (assuming homogeneous cell density 2*N/L*^2^)

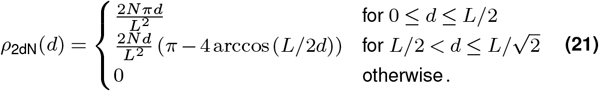

The cases in Eqn. (21) arise from the necessity to avoid ambiguity on a torus due to overlap. The connection probability *p*_2dN_(*d*) is small for *d* >500 μm, so we chose a layer size *L*=1 000 μm for which Eqn. 21 is well approximated by 2*Nπd/L*^2^ (Fig. M5a, solid vs. dashed line). The expected number of connected neurons is then given by 〈*k*_2dN_〉 ≈ 0.097*N*. To match 〈*k*_ring_〉=58 we would need *N*=600 cells in the 2d-network, which proved computationally difficult. We thus chose *N*=300 and increased the number of synapses *n*_syn,2dN_ per connected pair to compensate for the missing connections.

In order to constrain the conduction velocity *υ*_*c*,2dN_ and *n*_syn,2dN_, we matched the compound synchronous inhibitory conductance *G*_syn,2dN_(*t*) to the one expected for ring networks, see Eqn. (18) and Fig. M5d. Due to the linear increase in the number of cells at a given distance it follows that there are only few cells at very short distances, explaining the shift of *G*_syn,2dN_(*t*) towards later times compared to *G*_syn,ring_(*t*). We found that otherwise a good fit is established for *τ*_2dN_(*d*)=12s/m × *d* (i.e. a three times smaller conduction velocity *υ*_*c*,2dN_=0.0833 m/s than for ring networks) and *n*_syn,2dN_=7 (independent of distance, for sake of simplicity), see Fig. M5e. The respective delay distribution *f*(*τ*) is a rescaled version of *f*(*d*) with mean delay 〈*τ*_2dN_〉 ≈ 3.3 ms (shown for comparison in Fig. M5e).

**Figure M5.**
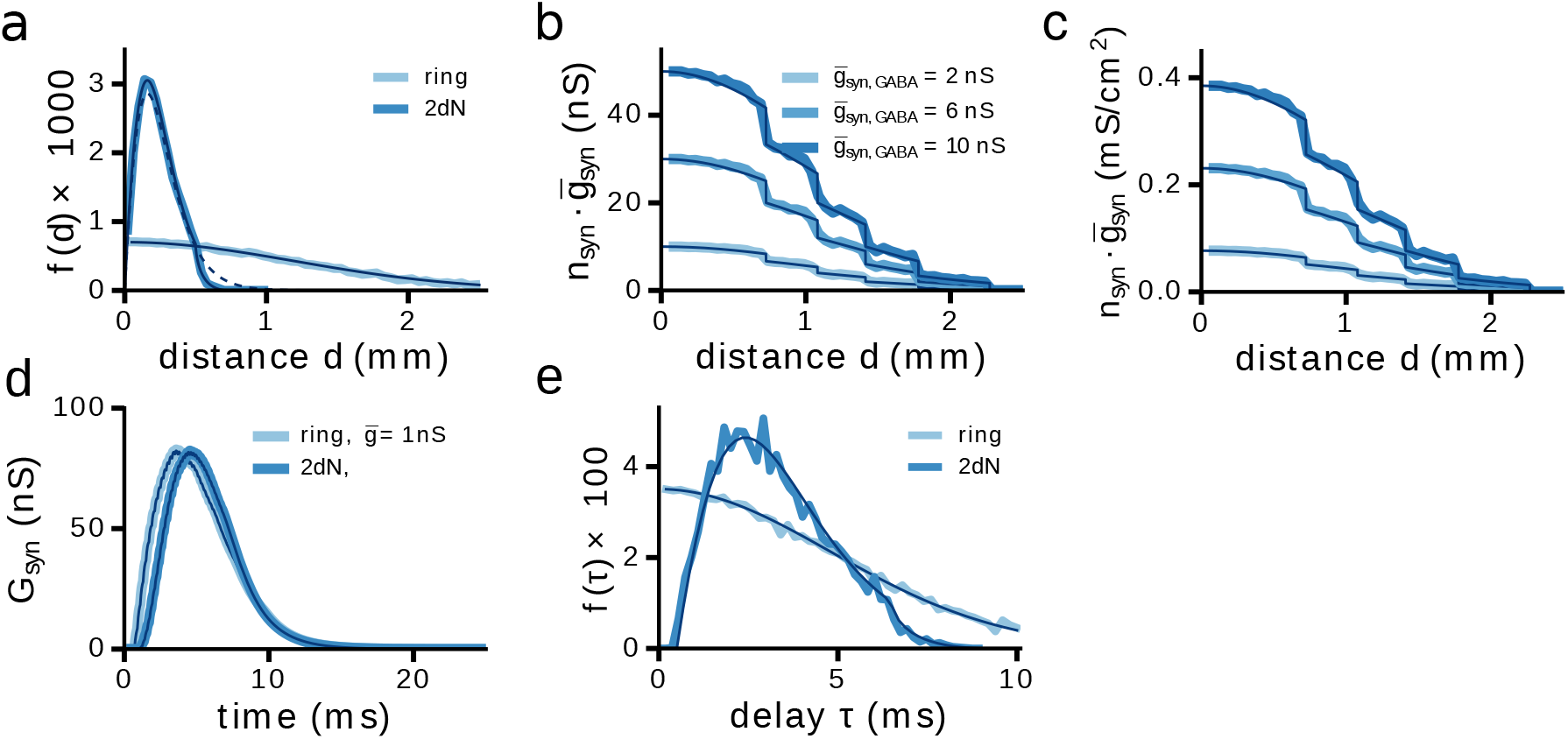
Connectivity statistics. **(a)** Density function of distances *f*(*d*) for the ring network (light blue) and the network embedded in two-dimensional space (darker blue) as estimated from a network instantiation versus the theoretical prediction (thin dark blue lines). For the two-dimensional (2dN) network we also show *f*(*d*) for the network without a maximal distance of *L*=1 mm (dashed line, see text). **(b)** Unitary, i.e., total inhibitory synaptic conductance between two PV cells at distance *d* for three different individual synaptic peak conductances 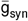. **(c)** Same as (b) in terms of the specific conductance, i.e., normalized by the cell area. Dark blue lines are respective analytical expectations. **(d)** Total inhibitory conductance received by a PV cell from all presynaptic cells assuming full synchrony for the ring network and peak conductance 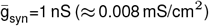 (light blue). The two-dimensional network was fitted to obtain a similar *G*_syn_ (dark blue, parameters *τ*_2dN_(*d*) = *τ*_0_ + 12s/m × *d* and *n*_syn,2dN_=7). **(e)** The respective delay distributions for ring and 2d-network after fitting *G*_syn,rand_. Wide lines in (d,e) are as measured from a network instantiations, dark blue lines are analytical expectations.

### Synchrony Index and Coherence

In several key studies (Bartos et al., 2002; Wang and Buzsáki, 1996; Bartos et al., 2001; Vida et al., 2006) a coherence-based measure was used to quantify synchrony in a network. In particular,

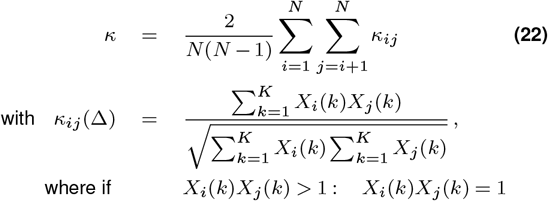

with binsize Δ, spike count histogram *X_i_* = hist[*S_i_*(*t*)|Δ] and number of bins 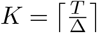 for considered time interval *T*. In the coherence measure employed in Bartos et al. (2002, 2001); Vida et al. (2006) the bin size is, moreover, adapted to the network firing rate *r* as Δ(*r*)=0.1/r. We find that the Synchrony Index FF/N as a global population activity-based measure is much simpler to quantify oscillatory behavior. The coherence measure potentially ignores spikes (it sets presence of any number of spikes per bin per neuron to one, see Eqn. (22)) and can also be high for non-oscillatory states, e.g., travelling waves. For the regimes we were considering, we found that the coherence measure *κ* also produces a floor at low or no synchrony (see Fig. M4b, M6d), which the FF-based Synchrony Index does not show. The FF on the other hand needs a very small time bin (smaller or equal to the sampling time) to correctly estimate FF=N for perfect synchrony (Fig. M6a), which for Poisson process driven neuron actvity is never quite the case. We optimized the time bin to yield maximal FF for a wide range of synchrony, see light blue solid curves in Fig. M6.

**Figure M6.**
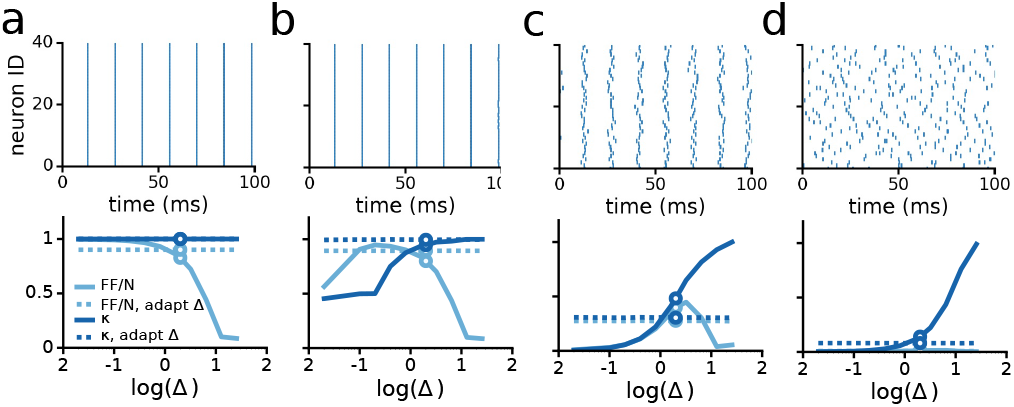
Fano-factor-based Synchrony index vs. Coherence Measure. **(a-d)** Upper row: Different variations of synchrony measures: Synchrony Index FF/N as function of fixed bin size (solid light blue), FF/N for adaptive bin size, i.e. dependent on the inverse of the firing rate (dashed light blue, firing rate is constant across panels at 70 Hz), coherence *κ* as function of bin size (Wang and Buzsáki (1996), solid dark blue, see Eqn. (22)), and *κ* for adaptive bin size (Bartos et al. (2002, 2001); Vida et al. (2006), dashed dark blue). Circle: δ=2ms is the value we consistently used to compute the synchrony index in our results. Lower row: respective spike data used to compute FF/N and *κ*. We used surrogate data, i.e., N=40 copies of the same regular spike train with increasing normally distributed jitter with standard deviation σ. (a) σ=0ms, (b) σ=0.1 ms, (c) σ=10ms, (d) σ=50ms.

## Supplementary information

**Figure S1.**
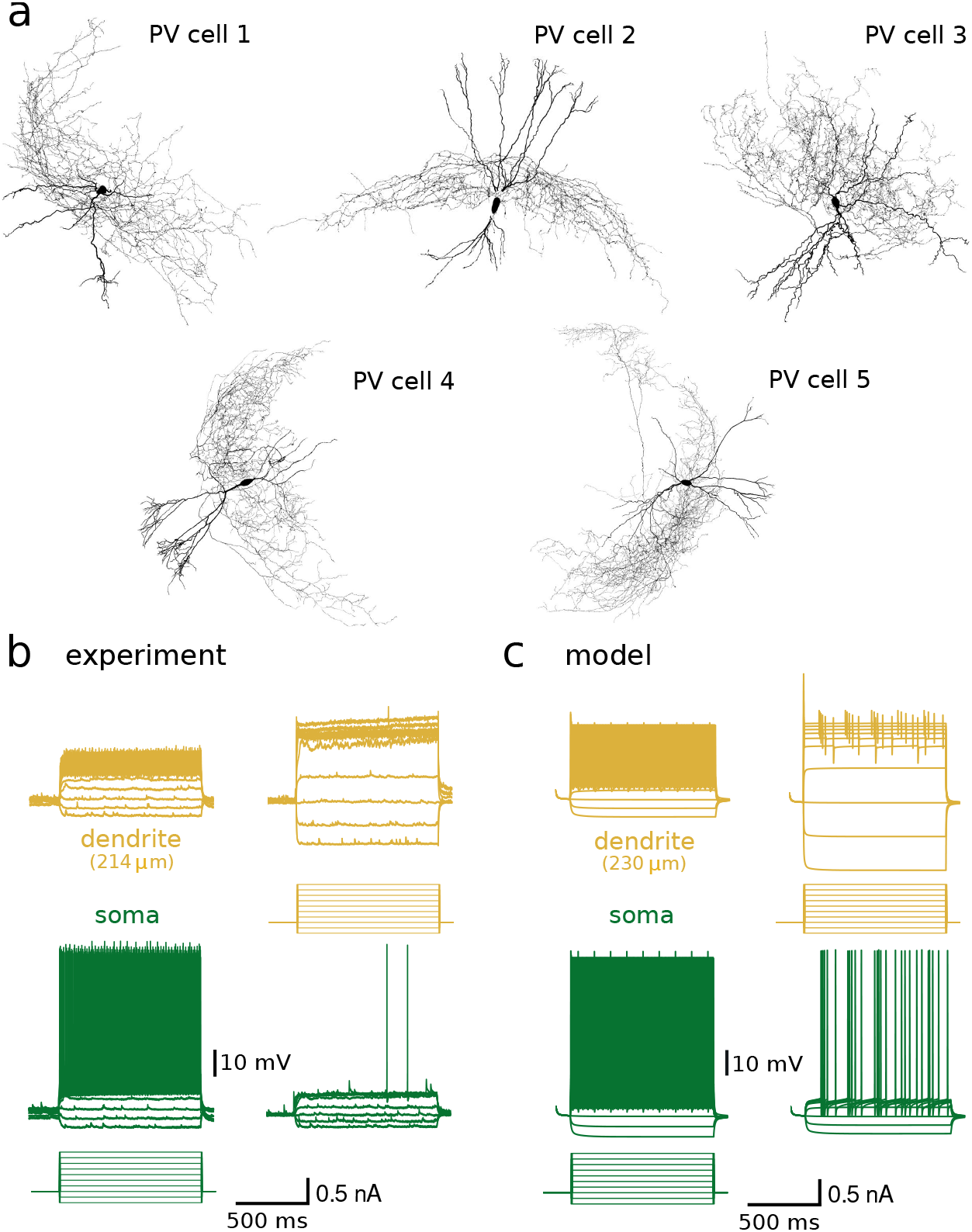
Morphology and physiological properties of the PV neuron models. **(a)** Reconstructed cell morphologies of five parvalbumin-expressing basket cells in the rat dentate gyrus used for simulations. **(b)** Example of dual soma-dendritic patch clamp recording. Voltage traces show soma and dendrite responses to current injections in either the soma (left) or the dendrite (right). Recording distance was 214 μm from the soma. **(c)** Same recording configuration as in (b). Model responses can be compared with experiments in (b). Model example is PV cell #2 (see (a), recording distance is 230 μm). Note that the fast transient at the onset of dendritic depolarization is a simulation artefact due to the instantaneous nature of the stimulus.

**Figure S2.**
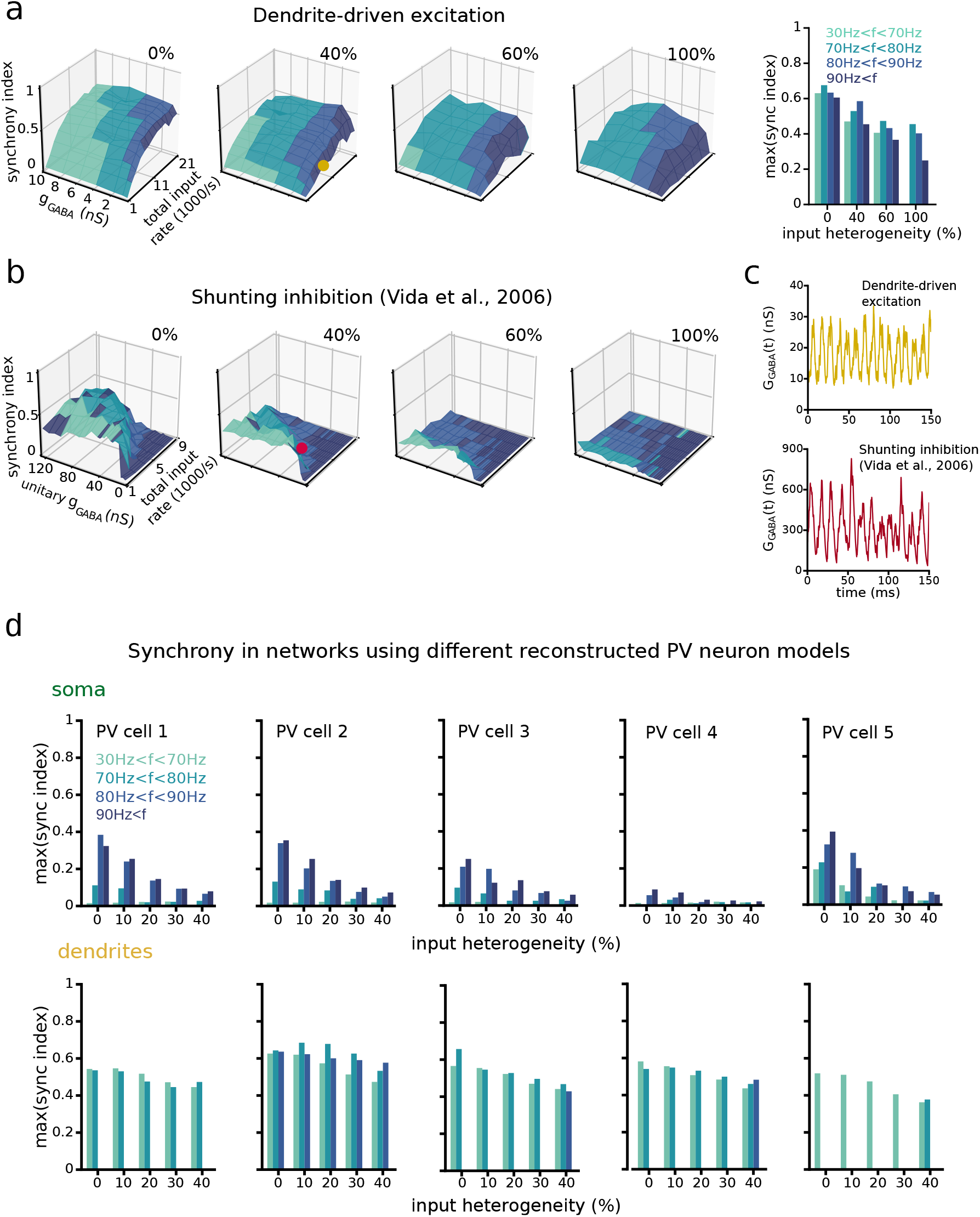
Robust dendrite-driven synchrony is maintained for high levels of input heterogeneity and is a property of all PV models tested. **(a)** 3D planes; Synchrony Index as a function of increasing input heterogeneity (0, 40, 60, 100%). Right, histogram of the maximum Synchrony Index per frequency band, as a function of input heterogeneity. **(b)** Synchrony Index of a previously published landmark model using single compartment cells and shunting inhibition (Vida et al., 2006). Same levels of heterogeneity as in (a). **(c)** Comparison of the total inhibitory synaptic conductance that is received by a single PV basket cell, using either the dendrite-driven model (top) or the the shunting inhibition model. The traces correspond to the orange and red dots on the 3D planes in (a) and (b). Note that we chose the network parameters with the lowest *g*_GABA_ that still produced synchrony. **(d)** Comparison of network simulations using the five different PV cell models. Histograms show the maximum Synchrony Index per frequency band, as a function of input heterogeneity.

**Figure S3.**
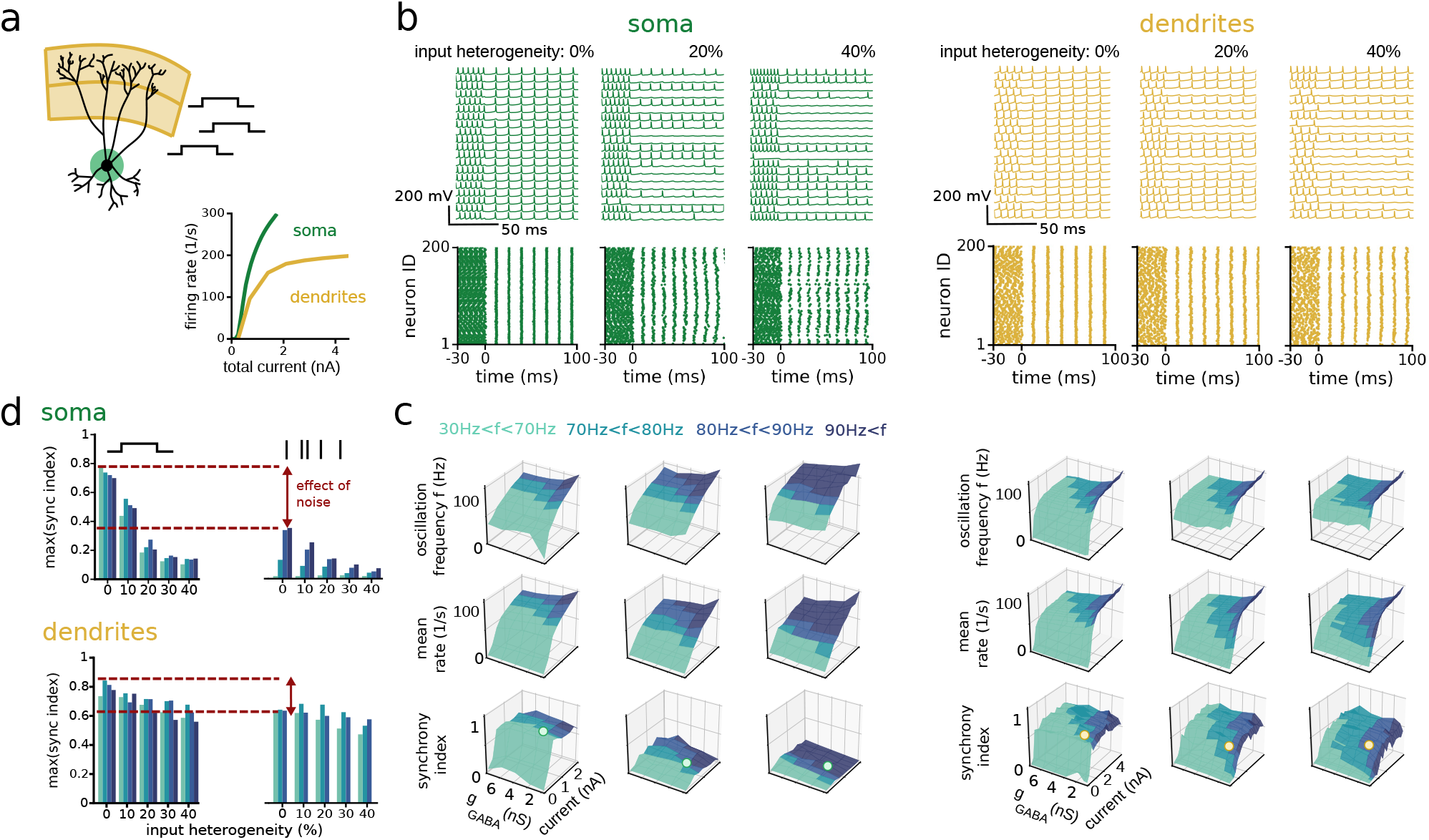
Noisy synaptic input is an important determinant of network synchrony. **(a)** Cartoon illustrating a PV neuron model driven by noiseless direct current (DC) inputs, distributed either perisomatically (green, 50 inputs ≤ 50 μm from the soma), or on the dendrites (orange, 100 inputs ≥ 120 μm from the soma). Bottom panel shows I-O relationships using either perisomatic or dendritic DC input. **(b)** PV cell activity in ring networks consisting of 200 PV neurons driven by perisomatic (green) or dendritic (orange) excitation. Input heterogeneity increases from left to right. Top row, example membrane potential traces showing spikes from 20 random cells. Bottom row, raster plots of all 200 PV cells in the network. Network starts uncoupled and inhibitory synapses activate at t=0 ms. **(c)** Network oscillation frequency (top row), average spike rate (middle row) and Synchrony Index (bottom row) as a function of total input rate and the strength of unitary inhibitory connections (*g*_GABA_). Colour legend shows the oscillation frequency ranges. The dots on the 3D planes of the Synchrony Index corresponds to the examples in (b). **(d)** The maximum Synchrony Index per frequency band, as a function of input heterogeneity for perisomatc and dendritic input (based on data in (b) and (c)). The Synchrony Index is shown for DC inputs (left panels) and for Poisson trains of synaptic conductances (right panels).

**Figure S4.**
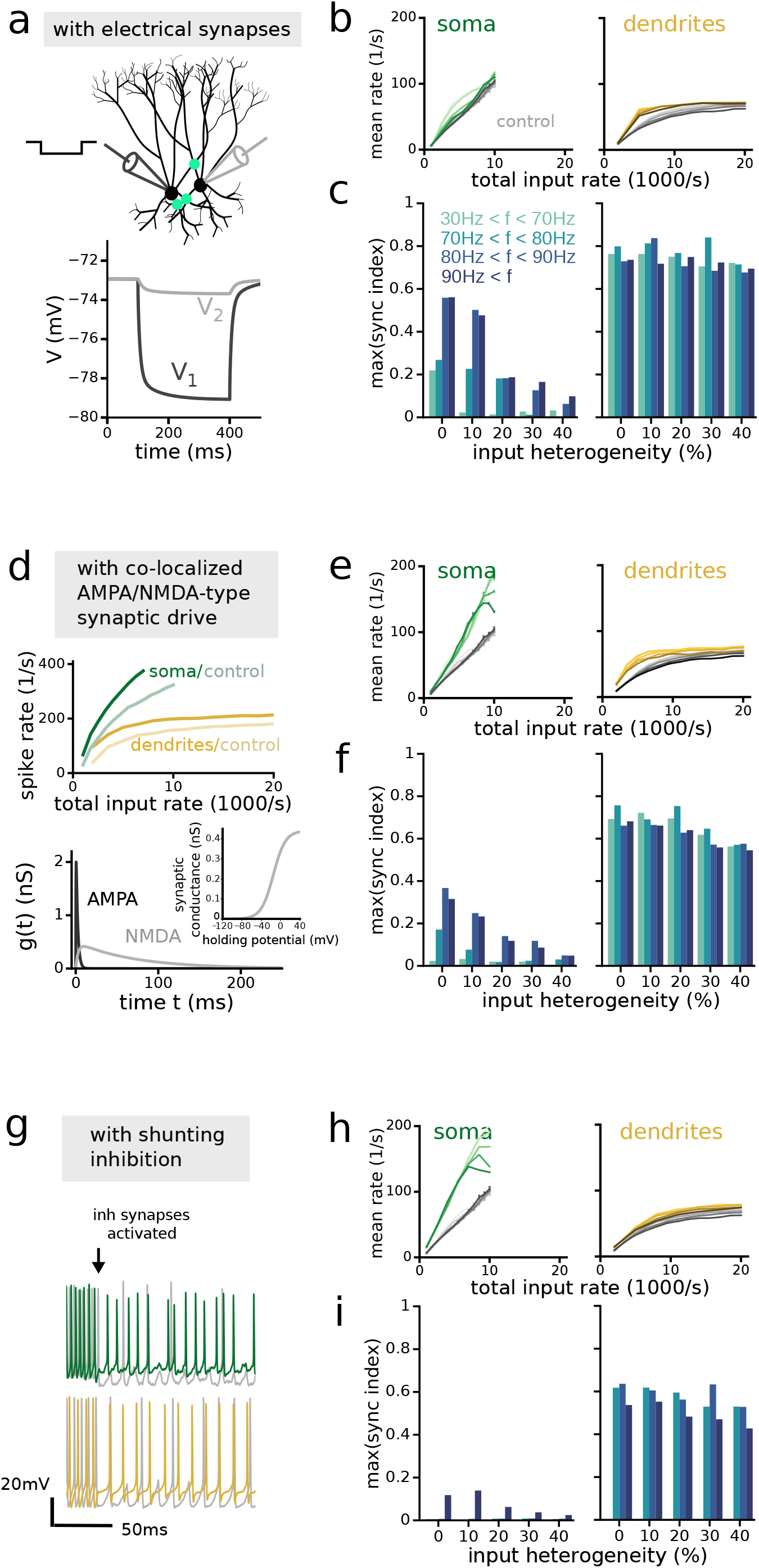
Enhanced robustness of dendrite-driven synchrony does not depend on electrical synapses, NMDA receptors or shunting inhibition. (a-c) Network simulations of PV neurons connected with electrical synapses in addition to chemical inhibitory synapses. (a) Cartoon of the recording configuration to test electrical coupling strength between two nearby PV neurons. Electrical synapses are inserted on the soma and proximal dendrites (≤ 50 μm) (see Methods Network models). Bottom, membrane potential responses of a coupled PV cell pair in response to a current pulse (−0.1 nA) in one of the cells. Electrical coupling strength is defined by the Coupling Coefficient (CC; the ratio of voltage changes in the cell pair, V2/V1 × 100). In this example, CC is 9%. (b) I-O relationships showing the mean spike rate of all PV cells in the network. This graph corresponds to a slice along the y-axis of the 3D plots as in Fig. 3c for *g*_GABA_=4 nS. Left, soma-driven networks. Right, dendrite-driven networks. Light to dark grey lines: increasing heterogeneity (0, 10, 20, 30, 40%) for networks without electrical synapses (control). Light to dark coloured lines, increasing heterogeneity for networks that also include electrical synapses. (c) The maximum Synchrony Index per frequency band, as a function of input heterogeneity. Left, soma-driven networks. Right, dendrite-driven networks. (d-f) Network simulations of PV neurons with co-localized AMPA and NMDA receptors. (d) Top, IO relationships of a PV neuron model, driven by excitatory input either close to the soma or on the dendrites. Faint curves are I-O relationships with only AMPA receptors. Bottom, kinetics and conductance of AMPA and NMDA receptors. The maximum conductance ratio between NMDA (0.42 nS) and AMPA receptors (2 nS) is ~ 0.21 (Koh et al., 1995b). Inset, voltage dependence of the NMDA receptors. (e,f) Same as (b,c) for networks with co-localized AMPA and NMDA receptors. (g-i) Network simulations of PV neurons with shunting inhibition. (g) Example voltage traces of PV neurons in a ring network with shunting inhibitory connections (*E*_rev_=−60 mV). Grey, simulation with hyperpolarizing inhibition. Green, soma-driven networks with shunting inhibition; Orange, dendrite-driven networks with shunting inhibition. (h,i) Same as (b,c) for networks with shunting inhibition.

**Figure S5.**
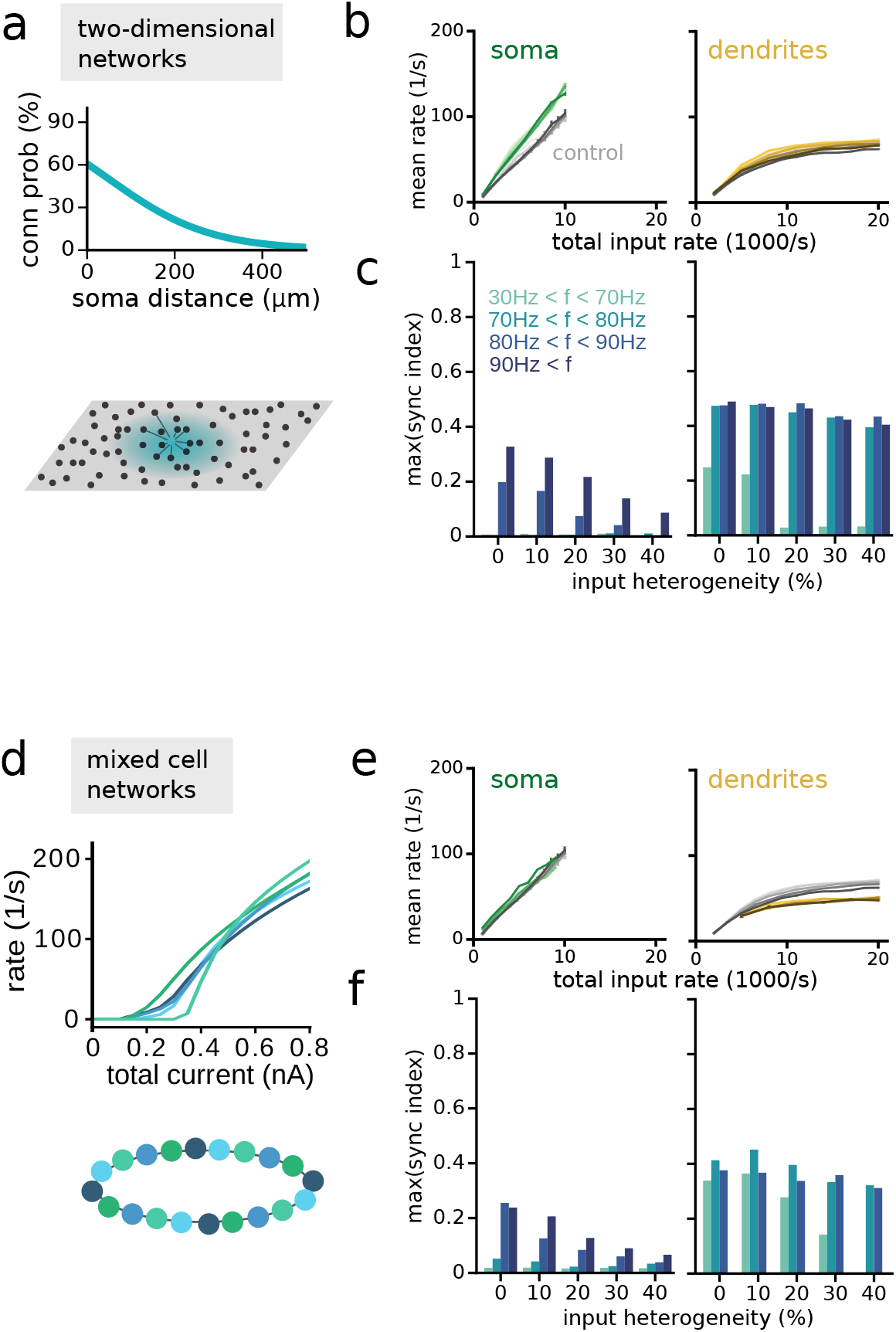
Enhanced robustness of dendrite-driven synchrony is maintained in PV neuron networks with a 2D organization and in mixed PV neuron networks. **a-c)** Simulations using PV cell networks with a 2D organization. PV cells were arranged on a 2D torus (see Methods). **(a)** Connection probability of PV cells as a function of distance between their somata. Adapted from experimental data on PV cells in the dentate gyrus reported in (Espinoza et al., 2018). Bottom, cartoon showing a 2D patch of PV neurons with local spatial connectivity. **(b)** I-O relationships showing the mean spike rate of all PV cells in the network (this graph corresponds to a slice along the y-axis of the 3D plots as in Fig. 3c for *g*_GABA_=4 nS). Left, somatic-driven networks (50 synapses ≤ 50μm from the soma). Right, dendrite-driven networks (100 synapses ≥ 120 μm from the soma). Light to dark grey lines: increasing heterogeneity (0, 10, 20, 30, 40%) for ring networks with local connectivity (control). Light to dark coloured lines, increasing heterogeneity for networks with random spatial connectivity. Most I-O relationships overlap. **(c)** The maximum Synchrony Index per frequency band as a function of input heterogeneity. Left, soma-driven networks. Right, dendrite-driven networks. **(d-f)** Simulations using networks consisting of a mix of all five PV models with a reconstructed morphology (see Methods). **(d)** Top, I-O relationships of all five PV models. Bottom, cartoon illustrating a ring network consisting of a mix of five different PV neuron models. **(e,f)** Same as (b,c) for mixed PV cell networks.

**Figure S6.**
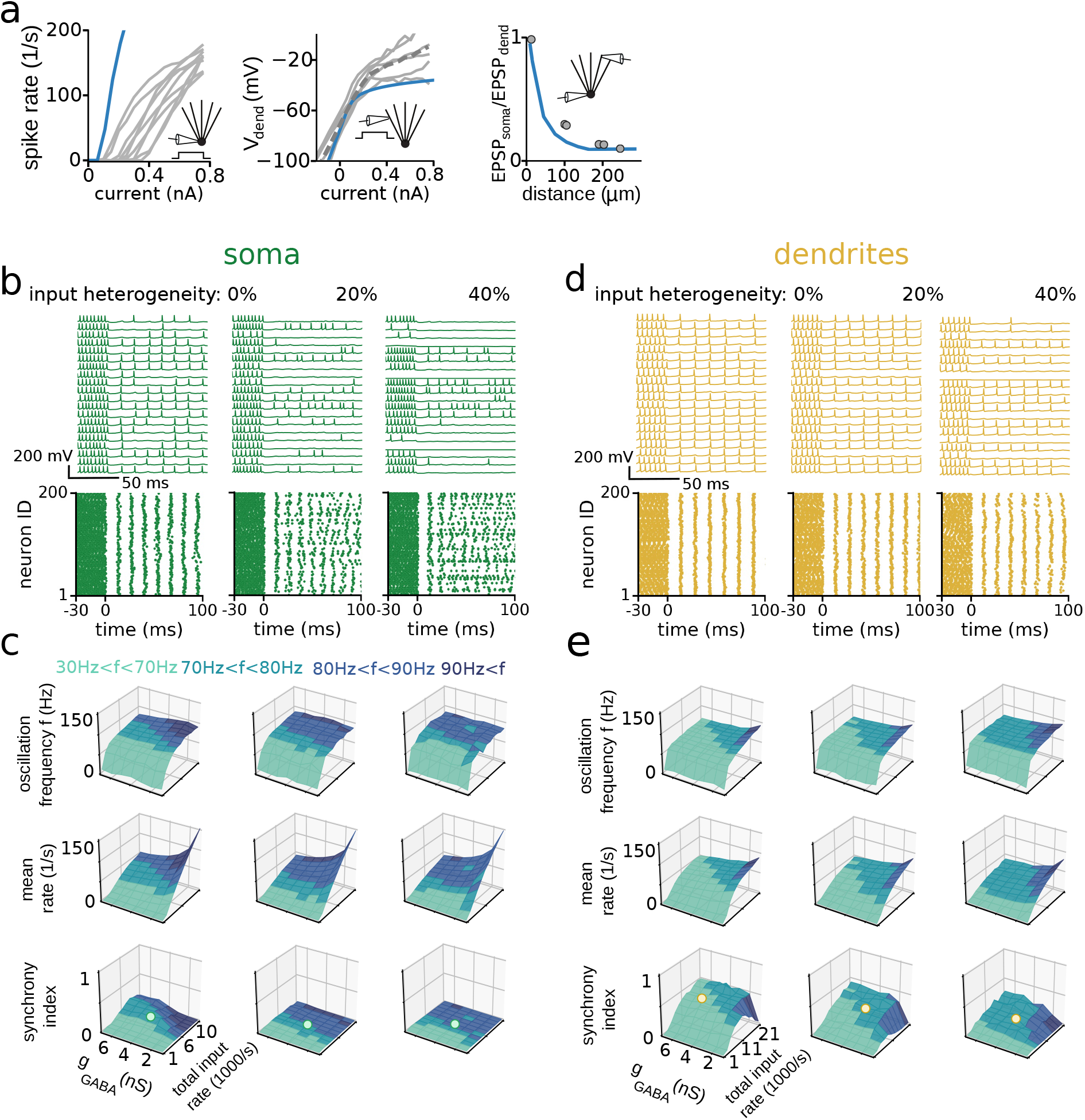
A simplified PV neuron model approximates the biophysical and network properties of the detailed PV models. **(a)** Comparison of the biophysical properties of the simplified model (blue) and the experimental data (grey, same experimental data as in Fig. 1 and 2). Left, I-O relationship when injecting current steps in the soma. Middle, I-O relationship when injecting current steps in the dendrite. Dendritic recording distance is 230μm from the soma. Right, attenuation of the EPSP amplitude during propagation from the dendrites to the soma. **(b)** PV cell activity in the network driven by perisomatic excitation. Input heterogeneity increases from left to right. Top, example membrane potential traces from 20 random PV cells. Bottom, raster plots of all 200 cells in the network. Network starts uncoupled and inhibitory synapses activate at t=0 ms. **(c)** Characterization of soma-driven networks as a function of input heterogeneity. Top, network oscillation frequency. Middle, average spike rate. Bottom, Synchrony Index. All are shown as a function of total input rate and inhibitory synaptic conductance. Colour legend indicates the oscillation frequency bands. White dots on the Synchrony Index corresponds to the data in (b). **(d)** As in (b), but for PV cell networks driven with dendritic input (100 synapses ≥ 150 μm from the soma). **(e)** As in (c), but for PV cell networks driven with dendritic input.

